# Protein reservoirs of seeds are composites of amyloid and amyloid-like structures facilitating sustained release during germination and seedling growth

**DOI:** 10.1101/2021.09.08.459376

**Authors:** Nabodita Sinha, Talat Zahra, Avinash Yashwant Gahane, Bandita Rout, Arnav Bhattacharya, Sangramjit Basu, Arunabha Chakrabarti, Ashwani Kumar Thakur

## Abstract

The seed protein functions and their localization in seed storage protein bodies (SSPB) are known for several decades. However, the structural and functional complexity of these SSPB is not known. Interestingly, the plant SSPB is morphologically similar to the amyloid-containing protein bodies found in other organisms and individual SSPB proteins were previously shown to form fibrillar structures under non-native conditions *in-vitro*. Therefore, we hypothesized that the seed storage protein bodies (SSPB) may have similar structures *in-vivo* for controlling seed functions. Since comprehensive *in-vivo* characterization of the SSPB and the structure-function relationship remains unexplored, we show firstly that wheat, barley, chickpea, and mungbean SSPB exhibit a speckled-pattern of amyloids interspersed in an amyloid-like matrix *in-situ*, suggesting their composite nature. This is confirmed by multiple amyloid-specific probes, biophysical characterization, electron-microscopy, peptide-fingerprinting, and differential degradation during germination. Moreover, the role of amyloid composites in seed germination is proved by the effect of signalling molecules and their correlation to germination parameters, using *in-situ* seed sections, *ex-vivo* protoplasts and *in-vitro* SSPB. These results would lay down foundation for understanding the amyloid composite structure during SSPB biogenesis and their structure-function evolution. It would further facilitate the exploration of molecular and atomic-level structural details of SSPB amyloids.

**Summary:** *Rationale:* The function of plant seed storage protein bodies (SSPB) in germination is known for decades. SSPB have aggregated and electron-rich morphology. However their structural complexity remains elusive. Based on their morphological similarity to amyloid-containing protein-bodies of other organisms, and amyloid formation by some plant proteins under non-native conditions, we hypothesized that SSPB might contain *in-vivo* amyloid structures for modulating seed functions.

*Methods:* To unambiguously identify seed amyloids in the presence of complex carbohydrate-structures of plant tissues, multi-spectral methods were used including amyloid-staining probes, high-resolution-transmission-electron-microscopy, x-ray diffraction and infra-red-spectroscopy. SSPB amyloid’s role in germination was shown using amyloid probes, MS/MS analysis, and plant hormones/proteases *in-situ* seed-sections and *ex-vivo* protoplasts.

*Key results:* The SSPB exhibit a composite structure of amyloid, amyloid-like aggregates and soluble proteins. During germination phases, the amyloids degrade slowly compared to the amyloid-like structures. Inhibition of amyloid degradation results in lower germination-index, confirming amyloid’s role in germination and seedling-growth.

*Conclusion:* The study for the first time illustrates the presence of composite amyloid structures *in-vivo* in plant seeds and determines their function in seed germination and seedling-growth. It would open original research questions for decrypting composite amyloid structure formation during SSPB biogenesis and their evolutionary advancement across plant species.

## Introduction

The original study of seed storage proteins in monocots and dicots can be traced back to the 18^th^ century.(Shewry *et al*., 1995; Shewry & Halford, 2002) The seed storage protein bodies (SSPB) of aleurone cells in cereals (monocots) are comprised of storage proteins including globulins,(Koziol *et al*., 2012) minerals and lipids.(Reyes *et al*., 2011; Isaienkov, 2014) In pulses (dicot), the cotyledon cells contain SSPB with globulins as the major storage proteins.(Derbyshire *et al*., 1976; Craig & Millerd, 1981) Most of these proteins are targeted to the SSPB in association with the Golgi apparatus, forming aggregated electron-rich structures.(Shewry & Halford, 2002) Similar structures are found in diverse species, ranging from bacteria to humans.(Schmidt, 2013) Interestingly, some of them also act as storage reservoirs for accumulating enzymes or proteasome units. (Narayanaswamy *et al*., 2009; Peters *et al*., 2015) These aggregates or inclusions are categorized as amyloids or amyloid-like structures to perform functions of storage and stability. Whereas amyloids are proteinaceous β-sheet-rich aggregates and exhibit apple-green birefringence with Congo red (CR), amyloid-like aggregates might lack one or more of the key characteristics of amyloids.(Benson *et al*., 2020; Matiiv *et al*., 2020) Although functional amyloids are well-studied in other organisms, there is prevailing confusion regarding plant amyloids. On one hand, groups have hypothesized that plants might not have amyloids owing to presence of anti-phenolic compounds in some plant tissues. (Surguchov *et al*., 2019) On the other hand, there are *in-silico* and *in-vitro* studies (Antonets & Nizhnikov, 2017; Antonets *et al*., 2020) regarding amyloid formation by individual plant proteins. But there is still lack of a detailed characterization according to current definition of *in-situ* amyloid detection.(Benson *et al*., 2020; Matiiv *et al*., 2020) Additionally, the role of the key molecular players in amyloid function is lacking. Since hormones and proteases play a major role in the maintenance/degradation of SSPB and subsequent germination, the effect of these molecular players on the amyloid aggregates need to be understood.(Guo & Ho, 2008) For this purpose, seed protoplasts are an ideal choice since the absence of cell walls reduces amyloid staining misperceptions. Also, protoplasts represent a dynamic active metabolic state and can be utilized to capture the effect of the exogenous effector molecules.

Although the protein content and types of the SSPB are known, their structural and functional complexity remains to be deciphered. Considering the structure-function relationship of protein bodies found in other organisms, we hypothesized that SSPB might possess amyloid or amyloid-like aggregates for performing seed physiological functions. In this study, we have used a combinatorial approach of multiple amyloid-specific probes and shown that the seed sections and protoplasts of wheat (*Triticum aestivum*) and barley (*Hordeum vulgare*), (monocotyledonous seeds with an endosperm as the major storage tissue) and chickpea (*Cicer arietinum*) and mungbean (*Vigna radiata*) (dicotyledonous seeds with two cotyledons as major storage tissue) exhibit a composite of amyloid and amyloid-like signatures in the SSPB of the aleurone and cotyledon cells. The amyloid-specific characteristics in the isolated SSPB and fibrillar nature of the amyloids shown by transmission electron microscopy (TEM)/high-resolution TEM (HRTEM), and x-ray diffraction, further bolsters the hypothesis. The amyloidogenic proteins are confirmed by laser capture microdissection followed by mass spectrometry. Moreover, the differential decrease of amyloid-like and amyloid structures during germination and seedling growth, and its correlation with specific germination parameters, confirms the role of SSPB amyloid composites in seed germination.

## Materials and Methods

a. **Materials:** Congo red (CR), thioflavin-T (ThT), acid fuchsin, 4′,6-diamidino-2-phenylindole (DAPI), papain, gibberellic acid (GA3), abscisic acid (ABA), phenyl methyl sulfonyl fluoride (PMSF), casein, ninhydrin, bovine serum albumin (BSA), sodium dodecyl sulphate, acrylamide, tetramethylethylenediamine, heptapeptide GNNQQNY and trypsin gold (mass spectrometry) were obtained from Sigma Aldrich. Proteostat^®^ aggregation assay kit was obtained from Enzo Life Sciences. Calcofluor white, xylene, ethanol, neutral buffered formalin (4%) and Pierce BCA kit were procured from Thermo Scientific. Paraplast for embedding was procured from Leica. Cellulase enzyme was obtained from SRL Chemicals. Dithiothreitol and iodoacetamide were procured from Merck. For desalination of the protein samples, Millipore C18 Ziptips were used. All other commonly used chemicals were obtained from Merck.
b. **Sectioning of the seeds and control samples:** Seed processing was done as per the standard protocols. (Wood *et al*., 2011; Jääskeläinen *et al*., 2013) Briefly, the seeds were halved and fixed in 4% neutral buffered formalin overnight. These were dehydrated with successive gradients of ethanol (30-100%) and permeated with paraplast to cut 8-12^μ^m sections (using Leica Microtome). The human fat biopsy tissue and potato tubers were processed similarly. Heptapeptide GNNQQNY fibrils were prepared in 1X PBS at a concentration of 2400 µM. The formed fibrils after 8 days were spotted on glass slides and stained. For **staining**, the slides were rehydrated and dipped in coplin jars containing the staining solution. Acid fuchsin (0.35%) and calcofluor white (10% v/v) were applied on the slides for 1 minute and DAPI (100 nM) was applied for 10 minutes in dark. Saturated solution of CR (80%) was applied for 20 minutes. Proteostat^®^ dye was used according to the manufacturer’s protocol. For ThT (acidified pH 4.5, 20μM), and Proteostat^®^ staining, 10 µL of the dye were added on the tissue sections or on slide spots in case of heptapeptide fibrils.(Navarro & Ventura, 2014) To ensure that the nuclei are not considered during quantification, we used a dual staining system of ThT and nucleus-specific DAPI. **(Figure S4 i1-i2)** Bright-field and fluorescence images were collected using Leica DM2500 fluorescent microscope equipped with cross-polarizers. ThT signal of seed sections was visualized by Leica TCS SP5 confocal system using He-Ne 488 laser (at 20% laser power) at 10X and 40X (under oil emersion).
c. **Isolation of protoplasts from wheat and mungbean:** The protoplasts from aleurone and cotyledon cells were isolated according to previously established protocols.(Taiz & Jones, 1971; Jacobsen *et al*., 1985) Briefly, 0.5 g wheat grains were de-embryonated, cut into quarter grains, and incubated in 10 mM arginine and 10 mM calcium chloride for 72 hours. After this, the endosperm was removed from the quarter grains under a dissection microscope to ensure protoplast isolation only from aleurone layer. The remaining tissue was incubated in cellulase solution (1 mg/ml, 0.3 units/mg) for 48 hours to free the protoplasts. For mungbean seeds, 0.5 g seeds were cut into 0.5-1 mm sections. These were incubated in cellulase solution (0.5 mg/ml, 0.3 units/mg) and swirled gently for 4-6 hours to free the protoplasts. The mungbean protoplasts show the typical structure of each cotyledon cell without cell wall and a cotyledon matrix with starch granules and SSPB. The wheat protoplasts of aleurone layer show a matrix devoid of starch, but SSPB presence is evident, as is typical for aleurone cells.
d. **Isolation and analysis of seed storage protein bodies (SSPB):** A protoplast count of 10000/ml (counted using hemocyometer) was used for SSPB isolation, and these were isolated using previously established protocols. (Bethke *et al*., 1996; Antonets *et al*., 2020) The protoplasts were added to double amount of lysis buffer containing 100 mM KCl, 2 mM MgCl2, 100 mM CaCl2, 50 mM sorbitol and 0.5% Triton-X at pH 7.2 in a chilled tube and incubated for 2 hours. The samples were layered on sucrose density gradients (20, 50 and 70%) and centrifuged at 37000g for one hour at 4°C. The SSPB were isolated from 50-70% layer (visualized the fractions and checked for SSPB presence). The isolated SSPB were centrifuged at 30000g for 30 minutes and resuspended in Tris-HCl (10 mM) buffer at pH 7.5. **For water-soluble albumin and salt-soluble globulin fraction isolation and physicochemical characterization of SSPB, please refer to Supplementary Section 1**.
e. **Germination conditions and parameters:** For checking germination parameters, each treatment was applied to 20 seeds (n=3). For detailed treatment, please see **Supplementary Section 1**. For germination, the seeds were first sterilized in 0.1% sodium hypochlorite, rinsed thrice in distilled water and placed on culture dishes with water or the exogenous molecules. The plates were sealed to reduce moisture evaporation and the seeds were incubated at 25±2°C, 12h dark/light photoperiod, 50±10% moisture growth chamber. The germination parameters including germination speed, rate and index were calculated. Germination speed is the average time after which 50% of the seeds have germinated. Germination rate is the percentage of seeds germinated whereas germination index is the ratio of germination percentage of treated seed vs. control. Germination was considered to be completed with 1 mm radicle protrusion. However, the experiment was conducted partially through post-germination phases as long as the seeds could be processed.
f. **Laser capture micro-dissection (LMD) and MS/MS analysis of the amyloid-containing tissues and protein fractions:** For LMD, 4-5 μm sections from mungbean and wheat (0 hour and 72 hours imbibition) were collected on glass slides. The 0-hour sections were stained with either CR or ThT and acid fuchsin. The 72-hour sections were stained with only CR to detect the changes in the amyloid proteins, as compared to 0-hour of imbibition (since by this time-point, the ThT-positive structures are almost not detectable). The earlier germination time-points were not considered, in order to capture significant changes in the type of amyloidogenic proteins. Leica LMD 7 was used to cut amyloid or amyloid-like regions (CR or ThT positive) and as negative controls, acid-fuchsin stained regions were cut. For sample cutting, both bright-field and fluorescence were used to demarcate the amyloid regions as is typically followed for amyloid diagnosis pipeline.(Nijholt *et al*., 2015) For each sample, 100,000 μm^2^ of area was cut to enable significant protein extraction (n=3). For processing of the LMD samples further for MS/MS, please refer to **Supplementary Section 1**.
g. **Computational analysis:** The secondary structure content of each protein was performed by UniProt, PDB, Pfam and JPred web servers. Aggrescan webserver was used to predict the potential amyloidogenic aggregation-prone regions in each of these sequences.
h. **Statistical and image analysis:** Origin Pro 9.1 was used to plot all the graphs and t-test was used to analyse the statistical significance between the data pairs. ImageJ and Leica Las X software were used to analyse the microscopy images. Each experiment is performed at least thrice (n=3) to obtain statistically significant results.

## Results

### Detection and confirmation of amyloid presence in seeds using combinatorial amyloid-specific probes

Before using amyloid-specific probes to detect amyloid presence in the seed storage protein bodies (SSPB), the seed tissue sections were dual stained with acid fuchsin and calcofluor white to visualize the proteinaceous and glucan-rich regions.(Burton & Fincher, 2014) It is essential to demarcate these two biochemical compartments since cellulose and β-glucans found in the cell walls of seed cells, also bind to Congo red (CR), a gold-standard dye for amyloids.(Herrera-Ubaldo & de Folter, 2018) Moreover, previously plant carbohydrates were referred to as amyloids due to their binding with iodine complexes and thus might result in nomenclature misperceptions.(Kooiman, 1960)

As shown in **Figure 1**, the seed coat, aleurone layer, subaleurone and endosperm cytoplasm of wheat and barley **(Figure 1 a1-a3)** and cotyledon cell-matrix of mungbean and chickpea seed sections **(Figure 1 b1-b3)** are stained with acid fuchsin and exhibit a pinkish-magenta colour on binding with proteins. Calcofluor white staining leads to a blue-fluorescence in the cell wall areas of barley **(Figure 1 a3)** and mungbean **(Figure 1 b3)** and represent glucan-rich regions. To further confirm the integrity and structural features of the seed sections, scanning electron microscopy (SEM) was performed on wheat and mungbean. In **Figure 1 c1-c2**, protein matrix of the aleurone and cotyledon cells is visibly intact, and corroborates with the structural features observed in the literature.(Kesari & Rangan, 2011)

**Figure 1.**
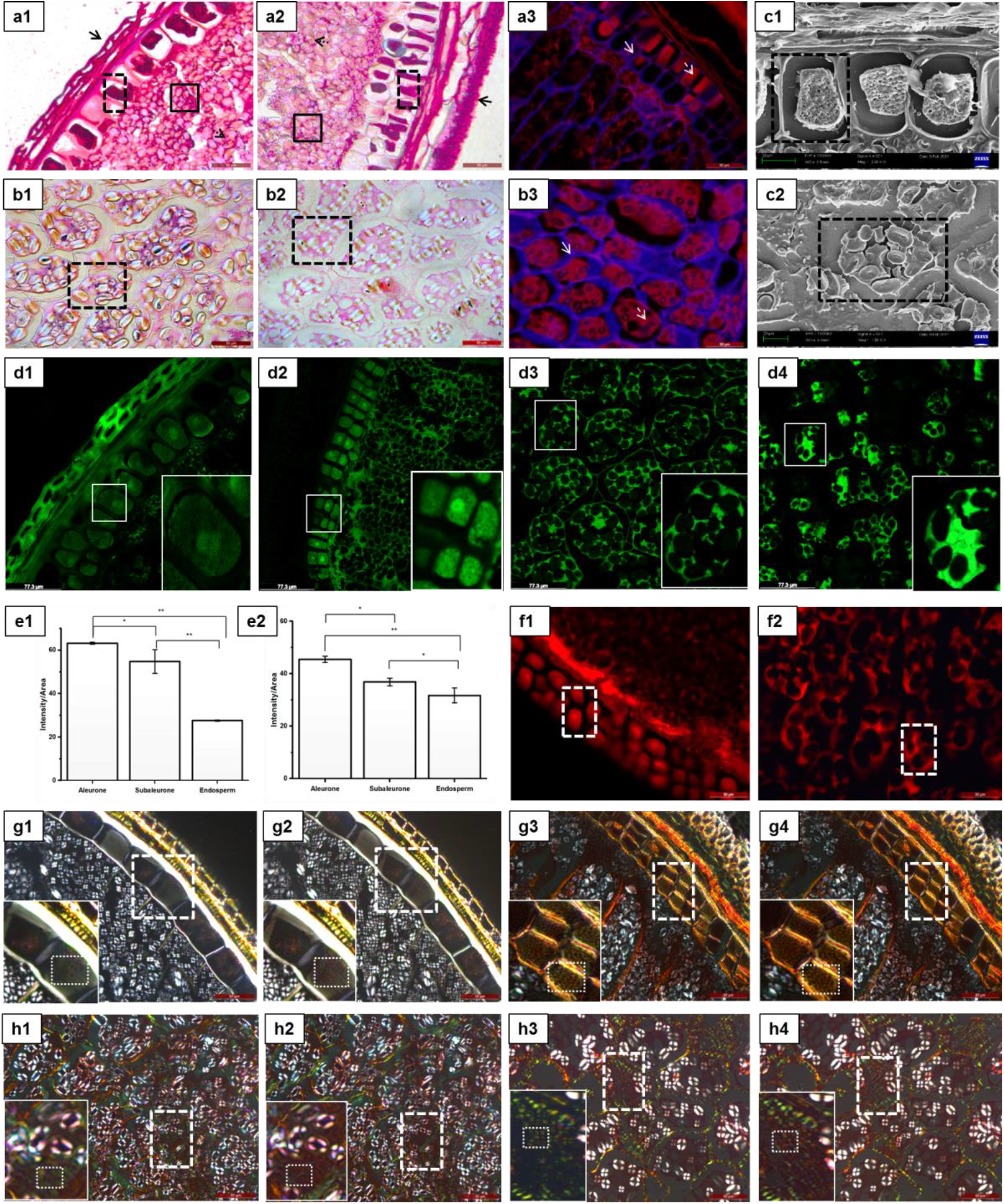
Analysis of amyloid structures in proteinaceous regions of seed sections. The protein-274 specific dye, acid fuchsin, exhibits characteristic magenta colour in the seed coat (solid black arrow), 275 aleurone (black dashed boxes), subaleurone cells (solid black box) and endosperm cells (black dashed 276 arrow) of wheat (*Triticum aestivum*) (a1) and barley (*Hordeum vulgare*) (a2). In the dicot seeds, the stain 277 is visible in the SSPB of cotyledon cells (black dashed boxes) of chickpea (*Cicer arietinum*) (b1) and 278 mungbean (*Vigna radiata*) (b2). In dual staining of acid fuchsin and calcofluor white, the barley aleurone 279 cells (a3) and mungbean cotyledon cells (b3) exhibit red fluorescence in proteinaceous regions, whereas 280 calcofluor white produces blue fluorescence in cell wall regions. Solid white arrows represent cell walls, 281 whereas dashed white arrows represent the SSPB. SEM (Scanning Electron Microscopy) analysis of 282 13wheat (c1) and mungbean (c2) reveal that the structure of the aleurone and cotyledon cells is maintained 283 after histological processing. The black dashed box represents an individual aleurone or cotyledon cell 284 with visible intact protein matrix. Gamma value for each acid fuchsin/calcofluor white image ranges from 285 0.6-0.7, the changes in brightness/contrast has been applied to the whole image. **ThT** staining of both 286 wheat (d1) and barley (d2) seed sections, exhibit an intense fluorescence in the aleurone layer, 287 suggesting an enrichment of amyloid-like protein aggregates in the aleurone layer. The dicot seeds of 288 chickpea (d3) and mungbean (d4) exhibit an intense ThT signal in the SSPB of cotyledons. The white 289 solid lined boxes represent fluorescing areas. The insets represent magnified portions of the solid-lined 290 boxes. The bar graphs represent each type of tissue’s intensity/area ratio, i.e., for aleurone, sub-aleurone 291 and endosperm in wheat (e1) and barley (e2).The ratio of five z-stacks are averaged for this purpose and 292 plotted as bar graphs with error bars representing standard error of the mean. Student’s t-test is 293 performed for statistical analysis. In (e1) *p=0.02, **p=0.001; (e2) *p=0.015, **p=0.003. Gamma and 294 intensity value for each confocal image is kept same for quantification purposes. **Proteostat**^®^ staining 295 exhibits that the monocot seeds of barley (f1) and dicot seeds of chickpea (f2) demonstrate the presence 296 of possible amyloids or amyloid-like aggregates in the SSPB of aleurone and cotyledon cells as evident 297 from the red fluorescence in these areas. White dashed boxes represent the intense signal in a 298 representative area. Gamma values for each ThT and Proteostat^®^ image range from 1.8-2.0, changes in 299 brightness/contrast have been applied to the whole image. **Confirmation of amyloids in seed sections** 300 **by CR staining**. The typical green-to-red birefringence when the sample is placed between two polarisers 301 at 40X magnification is shown. Both wheat (*Triticum aestivum*) (g1-g2) and barley (*Hordeum vulgare*) (g3-302 g4) aleurone cells show a visible change in the apple-green birefringence, characteristic of amyloids as 303 indicated by the white dashed boxes and the insets. Similar changes in birefringence are observed in 304 some regions of the SSPB of cotyledon cells of chickpea (h1-h2) (*Cicer arietinum*) and mungbean (*Vigna* 305 *radiata*) (h3-h4). Gamma values for each CR image are at 1.0. (Scale bars for acid fuchsin and calcofluor 306 stained images – 50 µm; for SEM images, scale bar – 20 µm; for ThT confocal scale bar – 77.3 µm; forProteostat^®^, scale bar – 50 µm; for CR staining, scale bar – 50 µm).

To demarcate amyloid or amyloid-like deposits in the seeds, we chose Thioflavin-T (ThT) as the first probe. It is one of the most popular dyes for amyloid or amyloid-like aggregate detection and studying *in-vitro* aggregation kinetics.(Biancalana & Koide, 2010; Boke *et al*., 2016) ThT dataset was validated by another molecular rotor, Proteostat^®^ to negate the possible artefacts resulting due to utilization of a single probe. It is a recently discovered probe, designed to detect intracellular and extracellular amyloids or amyloid-like deposits, even if the amyloidogenic proteins are sparse.(Oshinbolu *et al*., 2018; Laor *et al*., 2019)

For detecting amyloids or amyloid-like aggregates in seeds, wheat **(Figure 1 d1)**, barley **(Figure 1 d2)** chickpea **(Figure d3)** and mungbean **(Figure d4)** seed sections were stained with ThT and visualized using confocal microscope (10X images are presented in **Figure S1 a1-a4)**. In case of the wheat and barley seeds, intense fluorescence intensity was observed in the SSPB of protein matrix of the aleurone cells. The sub-aleurone layer shows intermediate fluorescence while the endosperm tissue shows non-significant ThT signal. The fluorescence intensity was quantified using ImageJ for each of these tissues to confirm this **(Figure 1 e1-e2)**. A uniform protein signal in the aleurone, subaleurone and the endosperm layer **(Figure 1 a1-a3)**, and significantly intense ThT signal in only aleurone cells of wheat and barley **(Figure S1 a1-a2)** signifies that although proteins are evident in all the tissues, amyloid and amyloid-like signals are prominently seen in the aleurone cells. In the mungbean and chickpea seed sections, the SSPB of cotyledon cell matrix produces an intense green fluorescence, suggesting presence of amyloid-like structures **(Figure 1 d3-d4; Figure S1 a3-a4)**. In Proteostat^®^ stained sections, the images show comparable pattern similar to ThT-results of barley and chickpea **(Figure 1 f1-f2)**, confirming the presence of amyloid or amyloid-like structures in SSPB.

However, *in-situ* staining with ThT is not enough as per the recent amyloid nomenclature (medical and functional) and detection guidelines. Most *in-situ* studies rely on CR as the gold standard probe for amyloid detection.(Benson *et al*., 2020) The optical anisotropy of amyloids on binding with CR and the resulting signature of apple-green birefringence, enables it as one of the most reliable methods for detection of amyloids in tissues and clinical samples.(Murphy *et al*., 2001; Benson *et al*., 2020) To confirm the presence of amyloids in the SSPB of aleurone and the cotyledon cells, seed sections were stained with CR and the samples were visualized between cross-polarizers.(Murphy *et al*., 2001; Benson *et al*., 2020) A characteristic apple-green birefringence of amyloids was observed in the aleurone SSPB of wheat **(Figure 1 g1-g2)** and barley **(Figure 1 g3-g4) (Movie S1)**. Fascinatingly, unlike ThT and Proteostat^®^ staining, which showed intense signal in the whole proteinaceous region of the aleurone cells, CR-induced birefringence was observed in some of these regions. Similar case was observed for the cotyledon cells of chickpea **(Figure 1 h1-h2) (Movie S2)** and mungbean **(Figure 1 h3-h4)**. The presence of CR-stained amyloids interspersed between amyloid-like structures suggests a composite structure of seed storage proteins.

To represent the inherent birefringence or autofluorescence of the tissue sections, the unstained sections were imaged using the polarizer and fluorescent microscopy, using same imaging parameters as that of stained sections and exhibited neither significant autofluorescence nor birefringence **(Figure S1 b1-b8)**. For positive control, human abdominal fat biopsy of a positive amyloid patient, **(Figure S1 c1-c2)** (Ghosh *et al*., 2021) and amyloid fibrils of Sup35-N terminal fragment heptapeptide GNNQQNY **(Figure S1 c3-c4)** were stained with ThT and Proteostat^®^ probes with same parameters as seed samples. Potato tubers **(Figure S1 c5-c6)** were chosen as the negative control, since these are rich in carbohydrates instead of proteins.(Shewry, 2003) Wheat endosperm tissue was used as another negative control, since it shows minimum signal with both probes **(Figure S1 c7-c8)**. ThT and Proteostat^®^ staining therefore suggest that wheat, barley, chickpea and mungbean seeds contain innate amyloid-like structures. As control of CR staining, the same tissue/fibrils used in ThT/Proteostat were used. **(Figure S1 d1-d8)**. To further corroborate amyloid-specific probe binding (fluorescence and birefringence), the glucan-rich cell walls of seed sections were digested with cellulase enzyme. Interestingly, although absence of Calcofluor white staining confirmed cell wall removal, CR-positive amyloid regions were evident. **(Figure S2 a-c)**. A control section without cellulase treatment (**Figure S2 d)** shows glucan-rich regions by Calcofluor white. Further, simultaneous staining of seed sections with Calcofluor white and ThT show non-overlapping spatial distribution of amyloids and glucans **(Figure S2 e-f)**. For unambiguous probe binding, the results are validated on protoplasts (cells without cell walls) and are discussed in the later sections.

### *In-silico* and biophysical characterization of seed storage protein bodies and confirmation of the amyloid and amyloid-like proteins using LMD-MS/MS

For finding the aggregation hotspots, web-servers such as Tango, Aggrescan and PASTA are generally used to assess amyloidogenic tendencies of proteins.(Belli *et al*., 2011) Aleurone and cotyledon cells contain globulins and albumins as the major storage proteins and are therefore analysed for their amyloidogenic potential. Globulins are well-characterized in terms of sequence and structure in comparison to albumins. **Figure S2 g-h** represents the secondary structure content of mungbean 8S (UniProt Q198W5), and soybean 11S globulin (UniProt P02858) respectively. These proteins belong to the cupin superfamily and are composed of β-barrel motifs. When the structural and sequence information of these proteins were analyzed by Aggrescan, interestingly, the high aggregation-prone regions were predicted mostly in the β-barrel structures, suggesting a plausible stacking of the barrels to form amyloids. A similar pattern was observed in other eukaryotic amyloidogenic proteins such as superoxide dismutase and bovine lactoglobulin **(Figure S2 i-j)**. As evident from the list in **Figure S2 k**, the globulin proteins of most cereals and pulses show an increased propensity of aggregation hotspots compared to the albumins, further strengthening that the SSPB globulins might be the major amyloidogenic proteins.

Since the SSPB is comprised of albumin and globulin as the major seed storage proteins, we isolated intact SSPB, and albumin, globulin fractions from SSPB **(Material and Methods d; Supplementary Section 1)**. When the SSPB of mungbean **(Figure 2 a)** and wheat **(Figure S3 a1)** were analysed by scanning electron microscopy (SEM), they show almost spherical structures. When the salt-soluble globulin protein fraction and SSPB of mungbean were analysed by dynamic light scattering (DLS), **(Figure S3 b1-b2)** the former shows a size of 2-30 nm whereas the SSPB show overall diameter of 400-600 nm and correlates with SEM analysis. Transmission electron microscopy (TEM) and high resolution TEM analysis of membrane-removed SSPB exhibit electron-rich morphology and fibrillar structures for both mungbean **(Figure 2 b1-b3)** as well as wheat **(Figure S3 a2-a4)**. ThT and CR staining of the SSPB of mungbean and wheat **(Figure S3 b3-b4)**, reveal green fluorescence and green-to-red birefringence respectively, characteristic of amyloid structures. The presence of ThT fluorescence in all the SSPB and CR-positive birefringence in some of these, further confirm the composite nature of the amyloids and amyloid-like assemblies.

**Figure 2.**
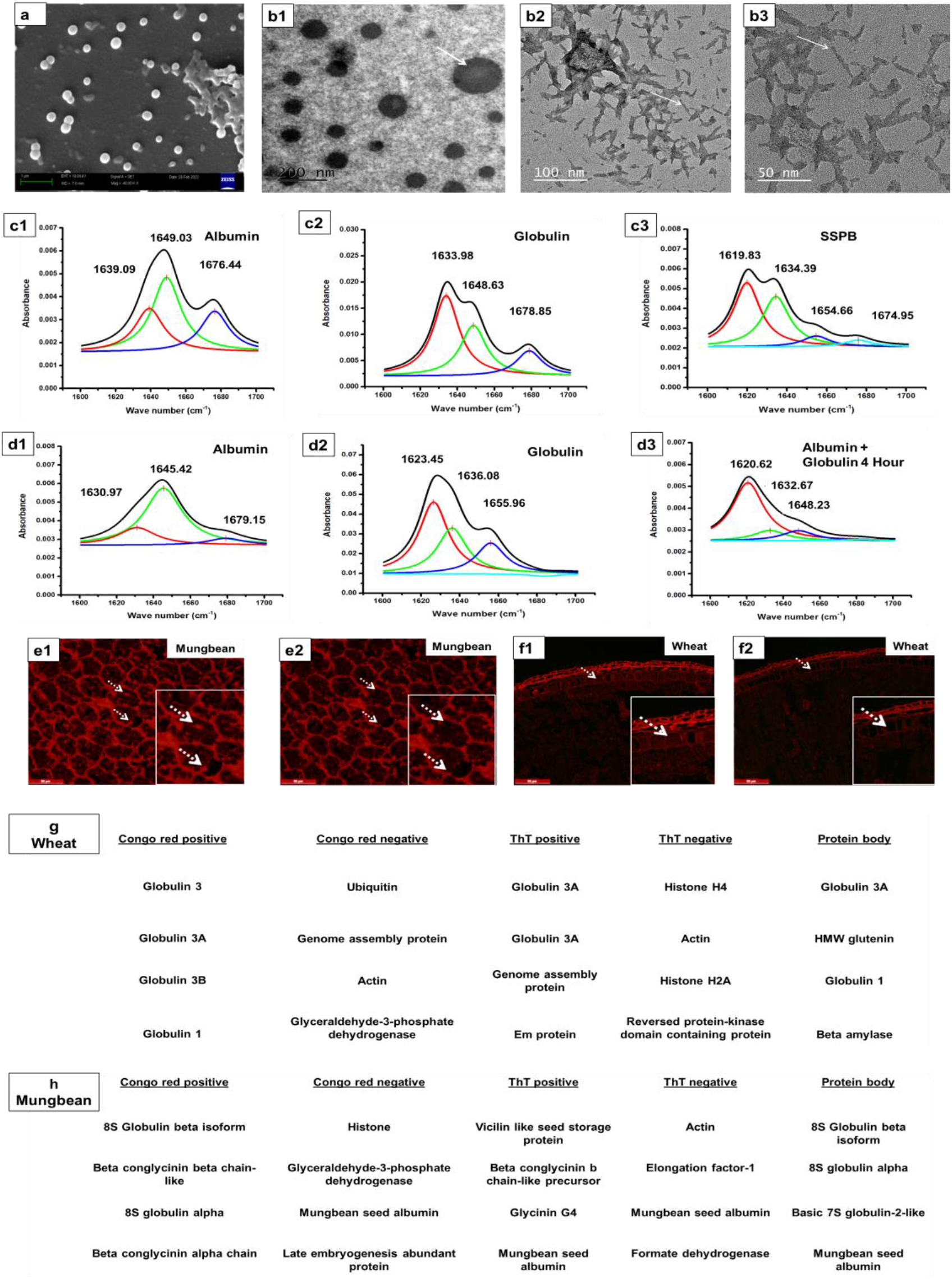
Analysis of the physicochemical properties of the seed storage protein bodies and 419 confirmation of amyloidogenic proteins: Representative SEM image (a) of isolated SSPB of 420 mungbean, reflecting on the globular shape and diameter as observed in literature. TEM (b1) and 421 HRTEM (b2-b3) analysis of the mungbean cotyledon SSPB shows electron-rich aggregates and fibrillar 422 structures upon removal of SSPB membrane. The white arrows point to the electron-rich structures of 423 SSPB and their fibrillar nature. FTIR analysis of the secondary structure content of water-soluble albumin, 424 salt-soluble globulin fraction and the SSPB of mungbean reveal that albumin consists of helical, sheet 425 and turn structures while globulins have predominant β-sheet signatures as evident by the major peak at 426 1633 cm-1. The SSPB on the other hand, shows evidence of amyloids by the shift towards 1619 cm-1(c1-427 c3). Dialyzed albumin of dicot mungbean shows the characteristic predominant helical signatures similar 428 to native protein isolated previously. Dialyzed globulins however show a shift towards 1623 cm-1 of the β-429 sheets, suggesting intersheet interactions of globulin at low salt concentration. On mixing and incubation 430 for 4 hours, the α-helix and the disordered structures decrease, while intersheet interactions become 431 more prominent, suggesting an increase in the overall amyloid signature (d1-d3). The LMD cut sections 432 before and after dissections are shown for mungbean (e1-e2) and wheat (f1-f2). The arrows represent the 433 CR-positive protein matrix cut regions. The inset boxes represent the magnified portions of the cut 434 regions. The proteins found after MS/MS of CR-positive amyloid areas, ThT-positive amyloid-like areas 435 and SSPB is represented in (g) and (h) (>95% confidence) (Scalebar of SEM -1 µm, TEM – 200 nm, HRTEM – 100 and 50 nm, LMD -50 µm).

Fourier-transformed infra-red spectroscopy (FTIR) can differentiate between the secondary structures of proteins and their aggregates, and often provide underlying differential signatures. In IR spectroscopy, the α-helix signature is found at ∼1654 cm^-1^, while β-sheets are mostly at ∼1635 and ∼1684 cm^-1^. The signature of the β-sheet rich amyloid fibrils are seen to be centered at ∼1610 cm^-1^ to 1632 cm^-1^ due to changes in the amide I band resulting out of interactions between β-sheets.(Waeytens *et al*., 2021) The turns and disordered structures are observed at 1670-1690 cm^-1^. Bovine serum albumin protein at same concentration was used as a control for the FTIR experiments.(Ahmad *et al*., 2016) The water-soluble albumin and salt-soluble globulin fractions isolated from SSPB, and the intact SSPB of mungbean **(Figure 2 c1-c3)** and wheat **(Figure S3 c1-c3)** were analyzed by FTIR. The albumin fraction isolated from the seeds, show predominant helical structure, (Moreno & Clemente, 2008) while the globulin fraction shows a predominant β-sheet structure at 1633-1636 cm^-1^ as expected from their β-barrel rich motifs.(Heyn *et al*., 2020) Interestingly, the SSPB not only shows the characteristic structural features of globulin, i.e. a predominant β-sheet richness, but also suggests inter-sheet interactions as observed by the shift of the bands towards 1616-1623 cm^-1^. Further, the helices and turn signatures are apparently evident, confirming the composite structure of the SSPB.(Sarroukh *et al*., 2013) The results suggest that the isolated proteins from SSPB in their soluble state exhibit their characteristic secondary structure signatures. However, when they assemble in the SSPB, they exhibit characteristics of amyloid and amyloid-like architecture.

Next, we wished to see whether the isolated soluble protein fractions could revert back to their amyloid state (the structures attained inside SSPB). To achieve this, the mungbean albumin and globulin proteins were dialyzed against buffer with reduced salt concentration. The albumin fraction, which is water soluble, **(Figure S3 d1-d2)**, shows no amyloid or amyloid-like signatures with ThT and CR after dialysis. However, the globulin fraction **(Figure S3 d3-d4)** and mixture of globulin and albumin **(Figure S3 d5-d6)** shows intense signatures of both ThT and CR. Interestingly, the same is reflected in the FTIR signatures of mungbean proteins. Dialyzed albumin shows α-helical and disordered structures while, dialyzed globulin shows a shift of the β-barrel structures towards 1623 cm-1, indicating amyloid formation. Further, a mixture of globulin and albumin (4 hours) show significant amyloid signatures and minor secondary structures (helices and turns) indicating a transition to amyloid composite form **(Figure 2 d1-d3)**. Therefore, based on amyloid-specific probe staining and FTIR studies, it is confirmed that, the globulin fraction or the mixture of globulin and albumin fractions can attain the amyloid composite state.

To further confirm the amyloidogenicity of SSPB fibrils and *in-vitro* reconstituted fibrils, these were analysed to check fibrillation kinetics, detergent resistance and x-ray diffraction signatures. The fibrillation kinetics of isolated and dialyzed globulin proteins of wheat and mungbean was monitored by ThT binding assay for different dialysis time-points. An increase in ThT fluorescence with increase in dialysis time, confirms the formation of amyloid structures **(Figure S4 a)**. These amyloid structures were analysed by TEM to check their fibrillar morphologies **(Figure S4 b-c)**. Next, the detergent resistance of the SSPB and isolated protein fraction fibrils were checked by SDS-PAGE. For this purpose, the disrupted or undisrupted SSPB fibrils, **(Figure S4 d)** and globulin fibrils **(Figure S4 e)** were boiled with SDS for different time-points (0-120 minutes). Without boiling, no bands for the samples are observed, suggesting that the constituent structures are unable to be resolved due to the large size of fibrillar aggregates. However, with increase in boiling time, more bands appear, suggesting that the fibrils are resistant to boiling in detergent for a particular time-point, but lose their detergent resistance after prolonged boiling. The control albumin (15-30 kDa) and globulin (30-60 kDa) isolated in their soluble form, (Quintieri *et al*., 2012; Yi-Shen *et al*., 2018; Kusumah *et al*., 2020) show their characteristic bands in gel. Powder X-ray diffraction of SSPB fibrils and isolated globulin aggregates was further performed to check the amyloidogenic signature reflections. In the SSPB fibrils of both wheat and mungbean, the equatorial and meridional reflections of amyloid structures are evident (diffused reflection at 1 nm and sharp reflection at 0.44 nm). Additionally α-helix (0.28 nm and 0.18 nm) and β-sheet (0.3 nm) specific reflections are seen, owing to the composite nature of the SSPB fibrils. For the isolated and dialyzed globulin fibrils of wheat and mungbean, primarily diffused amyloid-specific reflections were obtained. **(Figure S4 f)** (Eisenberg, 2003; Madine *et al*., 2008; Chakraborty *et al*., 2022) Together, the biophysical analysis confirms the amyloid nature of the SSPB fibrils and the ability of globulin proteins to attain the amyloid composite state.

Since the amyloidogenicity was indicated by the globulin protein fraction on dialysis, we wanted to confirm the major amyloidogenic proteins. For this, we performed laser capture microdissection (LMD) of CR positive, ThT positive and acid fuchsin positive fluorescent regions of wheat and mungbean sections. As representative examples, **Figure 2 e1 and f1** represents CR stained fluorescing areas of mungbean and wheat. **Figure 2 e2 and f2** represent the areas after dissection. This was followed by protein trypsinization and nano-LC-MS/MS for peptide fingerprinting of the dissected samples and the isolated SSPB. **Figure 2 g-h** represents the top four representative proteins predicted with highest confidence score (>95%) in case of wheat and mungbean. In both seeds, the CR-positive amyloid area is composed of mainly globulins. On the other hand, the ThT-positive area and SSPB in these seeds have globulins along with other proteins, suggesting the sequestration of soluble proteins inside the amyloid structures. To further confirm that the protein fractions isolated from SSPB, are actually the albumin and globulin fractions, these were also analyzed using MS/MS. **Table S1** shows the top scoring proteins of mungbean and wheat. The salt-soluble globulin fraction of both seeds is enriched in globulin proteins. The presence of seed storage albumin was evident in mungbean but not wheat, since albumin sequences of wheat are not available in UniProt using which the analysis was done. The number of peptides identified for each protein is represented in **Table S2**. The overall data confirms that globulins are the predominant amyloidogenic proteins in the SSPB and reflects on their composite nature. Further, the fibrillar nature of the amyloids in the SSPB is established, along with their amyloid-specific spectroscopic and diffraction properties.

### Functional role of amyloids during germination

Since amyloid-containing protein bodies in other organisms such as bacteria, perform functional roles, we hypothesized that SSPB amyloid composites might regulate seed physiological functions including germination.(Santos & Ventura, 2021) Classically, in *sensu stricto*, germination begins with water uptake by the seed (imbibition) and ends with the emergence of the embryonic axis, usually the radicle, through the structures surrounding it. To establish the role of SSPB amyloid composites during seed germination, wheat and mungbean seeds were imbibed in water and were monitored through the germination *sensu stricto* and post-germination phase, i.e. radicle elongation. At different time-intervals, the seeds were fixed, sectioned and stained with CR and ThT to detect the changes in the amyloids and amyloid-like structures. In the wheat seed sections, **(Figure 3 a1-a14)** the amyloid structures (CR-positive) were present till 48 hours but were not detectable at 72 and 96 hours in the aleurone cells, suggesting a possible degradation of these structures. In the mungbean seed sections **(Figure 3 b1-b10)**, the amyloids were detected upto 72 hours, suggesting that till these time-points, amyloids were not degraded to the full extent. For the dicot seeds, tissue processing was not possible beyond 72 hours due to fragile nature of the seeds. The representative seeds at each time-point are shown in **Figure 3**.

**Fig 3:**
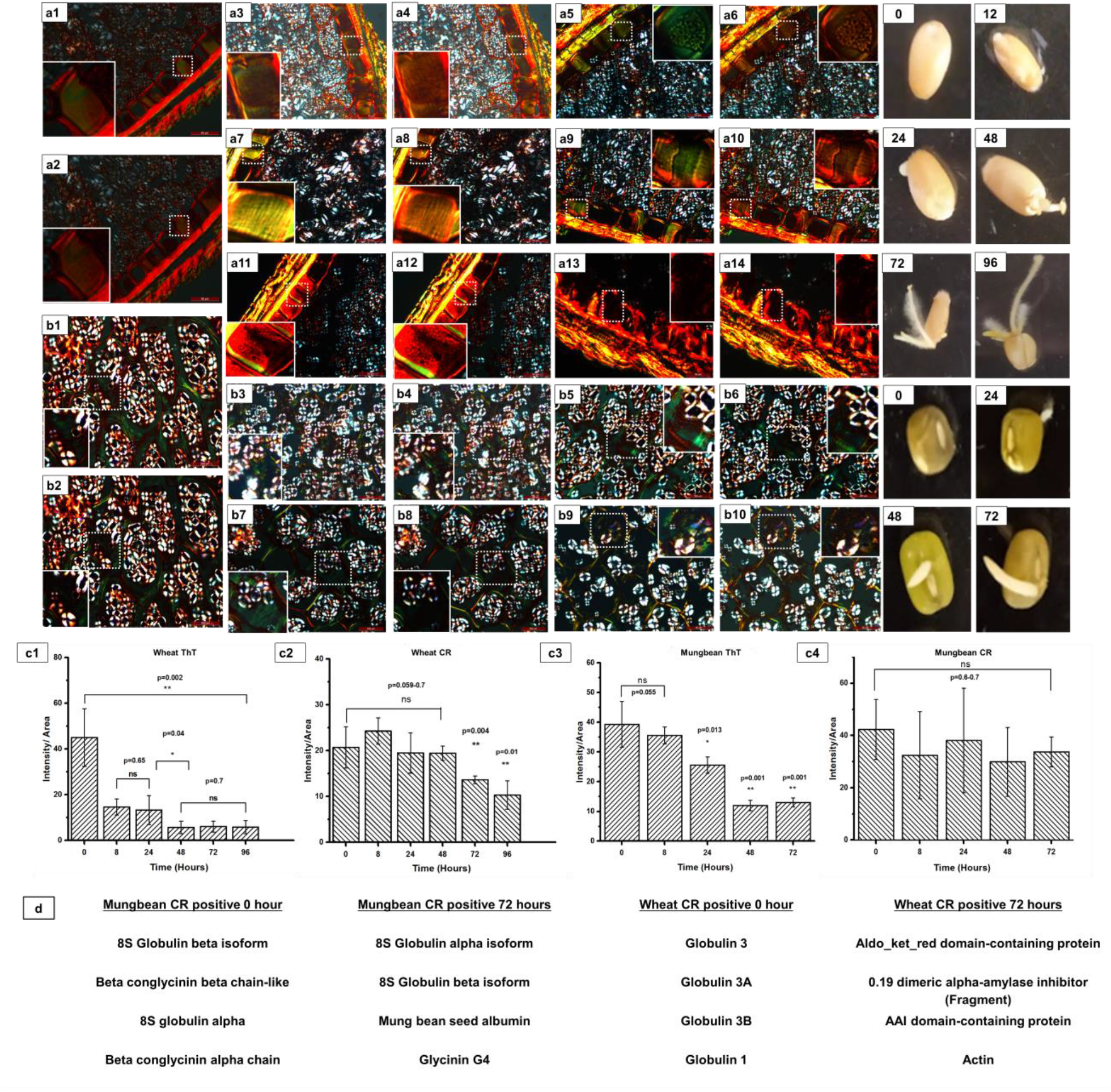
Congo red staining of germinated seeds, quantification of CR-ThT fluorescence and MS/MS 511 analysis of amyloidogenic proteins after germination. Congo red stained sections of wheat are 512 represented for germination at 0 (a1-a2), 4 (a3-a4), 8 (a5-a6), 24 (a7-a8), 48 (a9-a10), 72 (a11-a12) and 513 96 (a13-a14) hours, whereas the mungbean seeds were stained at 0 (b1-b2), 8 (b3-b4), 24 (b5-b6), 48 514 (b7-b8) and 72 (b9-b10) hours. In case of wheat, the amyloids are evident till 48 hours whereas in 515 mungbean, the amyloids are detected throughout all the time-points. White dashed boxes represent CR-516 positive representative areas and solid lined boxes represent the magnified portions of these areas. 517 (Gamma value for each image for CR staining is 1.3-1.6) Quantification of the signals are represented as 518 ThT of wheat (c1), CR of wheat (c2), ThT of mungbean (c3) and CR of mungbean (c4). ThT signal in 519 wheat start decreasing since 8 hours and in mungbean from 24 hours. The CR fluorescence however 520 decreases from 72 hours in wheat and does not decrease significantly till72 hours in mungbean. MS/MSanalysis of Congo red positive amyloid areas exhibit the decrease in the type of amyloidogenic globulins 522 after 72 hours of germination, suggesting their degradation in both wheat and mungbean (d).

In order to detect the changes in amyloid-like structures, the seed sections were stained with ThT after allowing germination and seedling growth upto 96 hours in wheat and 72 hours in mungbean. The ThT fluorescence of the seed sections was further compared to the CR fluorescence of the seed sections at the same time-points to quantify the changes in amyloid-like vs amyloid assemblies. The wheat seeds (**Figure S5 a1-a6 and Figure 3 c1)** show a decrease in ThT fluorescence intensity apparent from 8 hours while CR fluorescence decreases from 72 hours onwards **(Figure S5 b1-b6 and Figure 3 c2)**. Acid fuchsin staining of the germinated seed sections however depicts that the proteinaceous intensity decreases initially, but then remains stable throughout as shown in (**Figure S6 a-f and l1)**. In the mungbean seed sections **(Figure S5 c1-c5 and Figure 3 c3)**, the ThT fluorescence of SSPB shows a decrease in intensity from 24 hours, whereas the CR and acid fuchsin fluorescence remains similar till 72 hours as shown in **(Figure S5 d1-d5 and Figure 3 c4)** and (**Figure S6 g-k and l2)**. This shows that although less-stable amyloid-like structures are degrading, the more-stable amyloids degrade slowly and the overall protein content decreases first, and then becomes stationary **(Figure S7 a)**. The results suggest that the degradation of amyloids is accompanied or preceded by an upregulation of the soluble proteins including proteases and metabolic enzymes, thus maintaining the overall protein content (Han *et al*., 2013) and indicate towards a significant role of the amyloid degradation in germination.

To corroborate the properties of SSPB on similar lines, the isolated SSPB fibrils at each time-point of germination (0-72 hours) were analysed by SDS-PAGE. As evident in **Figure S7 b**, both wheat and mungbean SSPB at 0 hour, show no significant bands, suggesting the presence of fibrillar structures. However, with increase in time-points, the water-soluble albumin proteins (10-30kDa) and the salt-soluble globulin protein (43-71 kDa) bands become significant, suggesting their release from the fibrillar state. The results confirm that there is a step-wise release of proteins during germination.(Quintieri *et al*., 2012; Yi-Shen *et al*., 2018; Kusumah *et al*., 2020) The LMD samples of CR positive areas of wheat after 72 hours of germination show a drastic decrease in the types of globulin proteins, whereas the mungbean samples show significant globulin presence. This validates our germination data where wheat amyloids decline by 72 hours, but mungbean amyloids are evident till this time. **(Figure 3 d)**

SEM analysis of wheat **(Figure S5 e1-e2)** and mungbean seed **(Figure S5 f1-f2)** after 72 hours after imbibition shows a decrease in the overall SSPB content in the individual cells when compared to the 0 hour results. When the SSPB and globulin fraction isolated during different time-points for wheat and mungbean **(Figure S5 g1-g2)** are analysed for ThT, the globulin fraction shows baseline fluorescence. The SSPB however, show maximum fluorescence at 0 hours, which then decreases with germination, confirming loss of amyloid characteristics. When the same SSPB from wheat and mungbean **(Figure S5 h1-h2)** were analysed by FTIR, after 72 hours, the SSPB show a decrease in the relative predominance of the amyloid signatures and an increase in disordered structures. MS-MS analysis of the protein bodies after 48 h post-imbibition, also show a decrease in abundance of some globulin proteins. **(Figure S7 c)**

To further link the role of amyloid structures in germination, we incubated the water-soluble albumin, salt-soluble globulin, their dialyzed aggregated form and the SSPB with the purified seed endoproteases. (Supplementary Section 1) At each time-point, (0-8 hours) the dialyzed aggregated globulin fraction shows a slower release as compared to the soluble counterpart. On the other hand, SSPB shows the slowest release compared to the other fractions. To check whether the SSPB lipid membrane causes slower release, we removed the SSPB membrane and followed similar release assay. Despite membrane removal, the release remains significantly slow, confirming that the sustained release of peptides/amino acid is facilitated by the amyloid composites. **(Figure S5 i3-i4)**

The ThT-CR, MS/MS dataset and the *in-vitro* studies therefore suggest that the presence of the storage proteins in a composite amyloid structure facilitates sustained degradation during germination and seedling growth. The overall protein content however, does not decrease significantly suggesting that endogenous production of soluble proteins (enzymes, metabolic proteins) might be upregulated simultaneously.

### Linkage of amyloid degradation during germination with upregulation/inhibition of hormones and proteases

Germination in seeds is regulated by several plant hormones such as gibberellins (GA) and abscisic acids (ABA). GA further acts downstream to regulate protease expression after seed imbibition and therefore acts as one of the triggers of germination. ABA, on the other hand, acts as an inhibitor of GA to maintain dormancy.(Shu *et al*., 2018) The initial seed proteases are present inside the SSPB, and can cleave the disulphide bonds of the globulin proteins and prepare them for further cleavage. The subsequent proteases are expressed in the cotyledon or aleurone cell cytoplasm and are assumed to cleave the storage proteins. These endoproteases are primarily of cysteine or serine protease type and can be inhibited by protease inhibitors including phenyl methyl sulfonyl fluoride (PMSF).(Otegui *et al*., 2006) Although, GA leads to a degradation of the seed storage proteins, the underlying mechanism of the degradation control is not deciphered in details previously. Thus hormones and protease-dependent amyloid degradation might aid in controlled germination and post-germination phases.

To gain an insight into how the amyloids are degraded in response to the hormones and proteases, the wheat and mungbean seeds were imbibed in water, GA, ABA, GA and PMSF, ABA and PMSF, and PMSF alone for 72 hours. **(Supplementary Section 1)** It was observed that, as compared to water (control), **(Figure 4 a1-a4, g-h)**, the seeds incubated in GA alone, **(Figure 4 b1-b4, g-h)** showed decrease in amyloid signal. On the other hand, GA and PMSF treatment gave stronger amyloid signal, suggesting that amyloid degradation rate is further decreased in presence of PMSF as compared to normal germination in water. **(Figure 4 c1-c4, g-h)** ABA **(Figure 4 d1-d4, g-h)** and PMSF **(Figure 4 f1-f4, g-h)** imbibition when used individually, show increased amyloid and amyloid-like content when compared to control, indicating that the amyloid degradation is inhibited more in these cases. Interestingly, a simultaneous imbibition with ABA and PMSF treatment, shows maximum amyloid signatures among all treatments **(Figure 4 e1-e4, g-h)**. On analysis of the germination parameters, (germination speed and germination index), it was observed that the decrease in amyloid signal in case of GA treatment is accompanied with a faster germination and higher germination index, while adding PMSF alongside GA counters the effect. Interestingly, the increase/maintenance in amyloid signal (ABA, PMSF individually and together) correlates with a significantly lower germination speed and germination index **(Figure 4 i)**. Representative seeds and their radicle length after 24 hours of imbibition is shown in **Figure 4 j**. Overall, the CR and ThT data and the correlation of amyloid presence with germination parameters, suggests that the degradation of the amyloid and amyloid-like composites, plays an important role in controlling germination and seedling growth.

**Figure 4:**
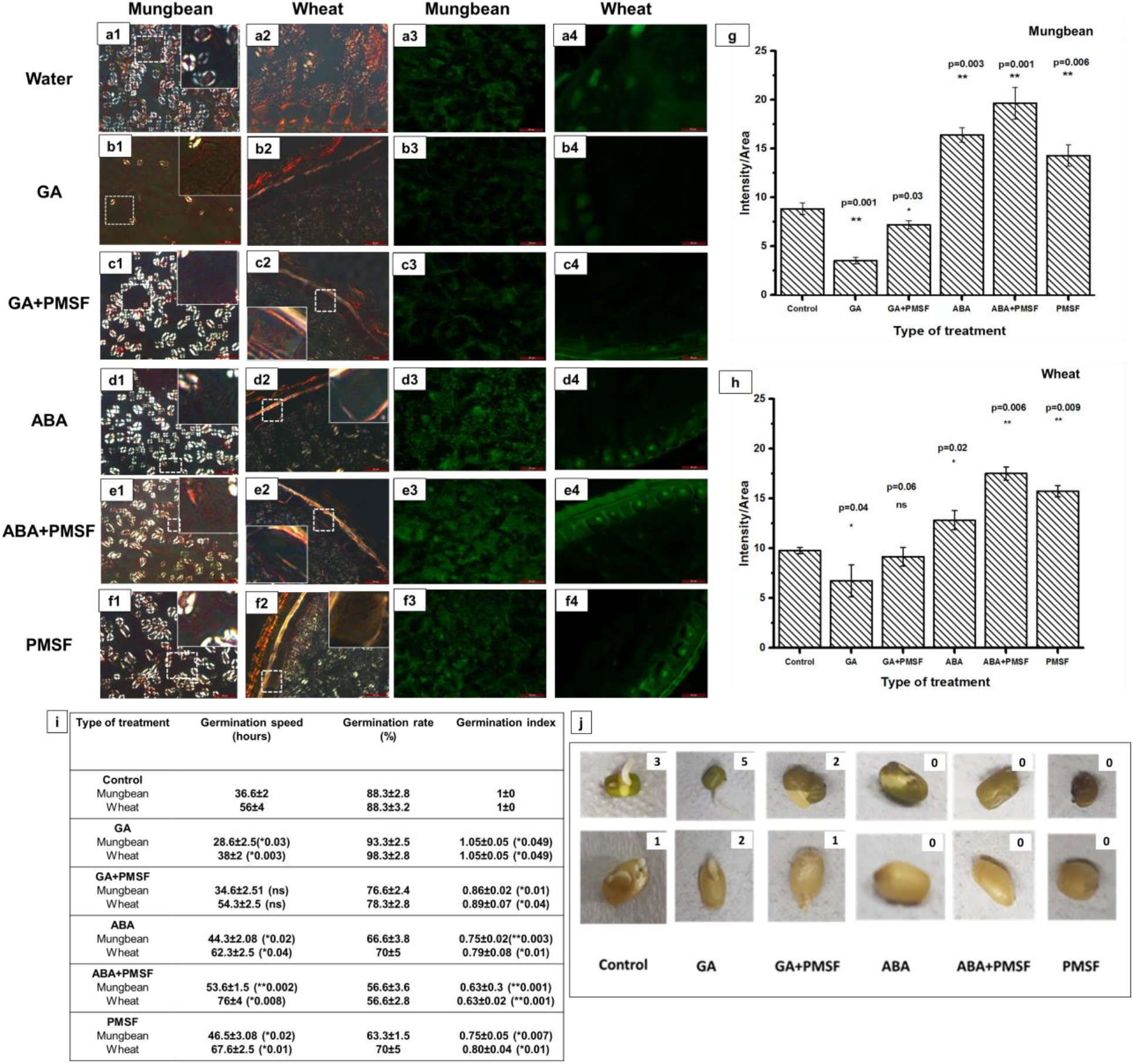
Exogenous treatment of wheat and mungbean seeds with hormones, proteases and 560 inhibitors and their link to amyoid degradation during germination. The mungbean and wheat seeds 561 were imbibed with water (a1-a4), GA (b1-b4), GA and PMSF (c1-c4), ABA (d1-d4), ABA and PMSF (e1-562 e4), PMSF (f1-f4). The left two columns represent the CR-stained sections of mungbean (a1-f1) followed by wheat (a2-f2). The white dashed boxes represent the insets and the solid white boxes represent the 564 magnified portions of the insets. The right two columns represent ThT stained sections of mungbean (a3-565 f3) followed by wheat (a4-f4). The ThT signature intensity of the seed sections was further quantified 566 using ImageJ and plotted (g and h), for mungbean and wheat seed respectively. Bars represent mean 567 values with error bars as standard error of mean. (Gamma values range from 0.8-1.0, Scale bars for each 568 image corresponds to 50 µm). The germination parameters are recorded (i) for 20 seeds total in each 569 group and the data is average of three such replicates. The radicle length and the representative images 570 of the seeds after treatment are shown in (j) after 24 hours of germination. The insets represent the 571 radicle length in mm. The upper panel shows the mungbean seeds whereas; the lower panel shows the wheat seeds.

To check the amyloid signals in the physiological context and in a viable state of the cells, we isolated and used protoplasts. These have an active metabolic state and can be investigated for the real-time effect of exogenous molecules (Jacobsen *et al*., 1985) on the amyloid content. Further, due to absence of cell walls, there is minimum chance of interference in staining. Since, protoplasts do not require dehydration, fixation or embedding in resin, it negates out the plausible artefacts caused by tissue processing methods. At first, the presence of amyloids in the SSPB of the wheat and mungbean protoplasts were shown using ThT **(Figure 5 a1,c1)** and CR **(Figure S9 a1**,**a5)**. In both wheat and mungbean, the intense ThT fluorescence and green-to-red birefringence characteristic of amyloids are apparent. To confirm the tissue origin of the protoplasts, the dissected aleurone cells and for staining control, leaf protoplasts with ThT staining are shown. **(Figure S9 f1-f4)**

**Figure 5:**
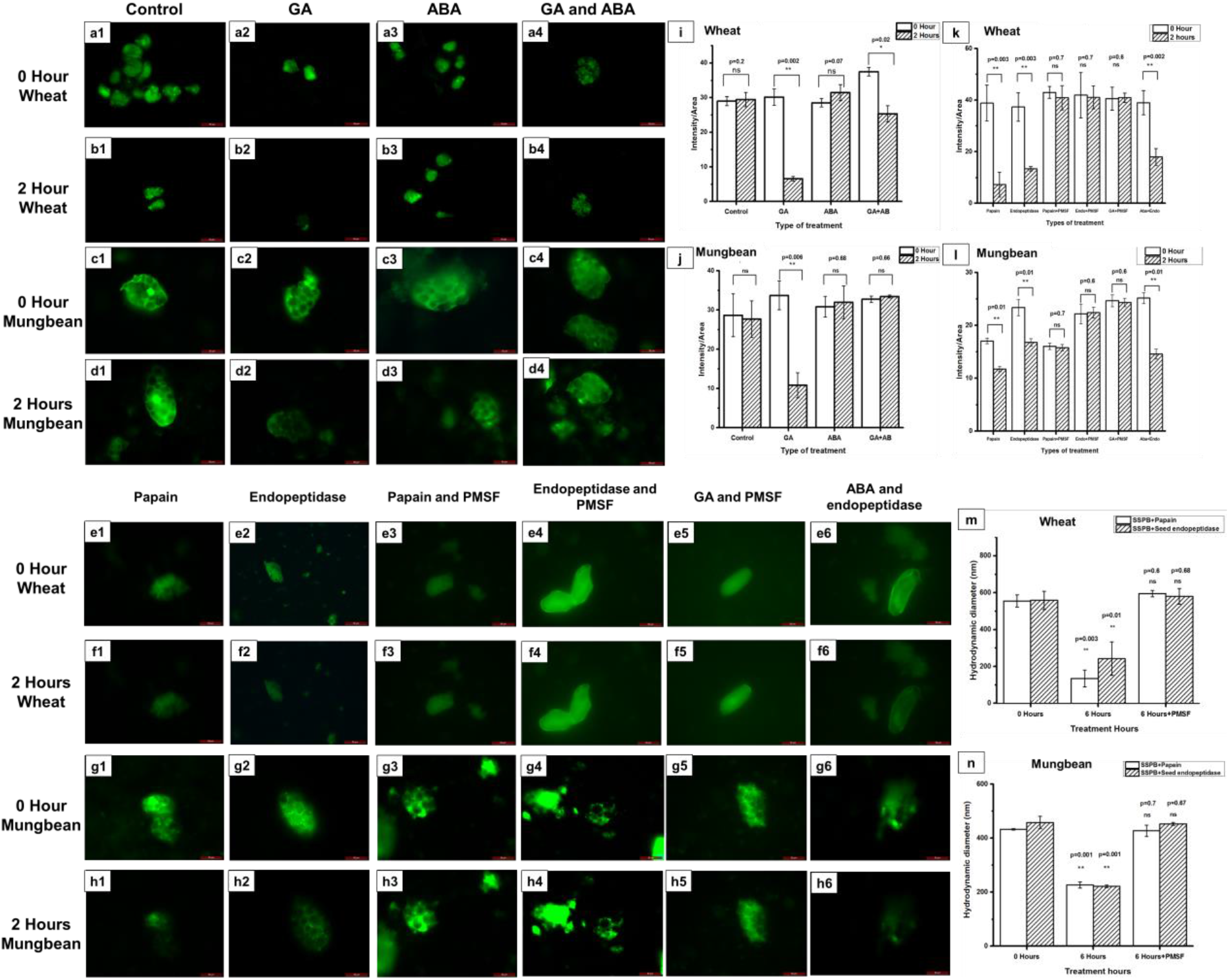
Effect of plant hormones and proteases on amyloid aggregates. Wheat aleurone 631 protoplasts treated with buffer, gibberellin (GA), abscissic acid (ABA), GA and ABA at 0 hour (a1-a4). The 632 protoplasts were treated for 2 hours with the same treatment (b1-b4). 0-hour mungbean cotyledon 633 protoplasts treated with buffer, GA, ABA, GA and ABA, (c1-c4) whereas protoplasts after similar treatment 634 of 2 hours are shown in (d1-d4). GA treatment decreases the amyloid signal whereas ABA treatment 635 leads to retention of fluorescence. A combined treatment of these hormones, show decrease in 636 fluorescence in the wheat but not in mungbean protoplasts. The quantification of the ThT fluorescence is 637 analysed by ImageJ and represented as (i) for wheat and (j) for mungbean. Wheat (e1-f6) and mungbean 638 (g1-h6) protoplasts were treated with papain (e1,f1,g1,h1) endopeptidase (e2,f2,g2,h2), papain and 639 PMSF (e3,f3,g3,h3), endopeptidase and PMSF (e4,f4,g4,h4), GA and PMSF (e5,f5,g5,h5), ABA and 640 endopeptidase (e6,f6,g6,h6) at 0 hours and after 2 hours. The signatures were quantified and plotted in 641 (k) and (l) for wheat and mungbean respectively. Treatment with the proteases decrease the amyloid 642 signatures, whereas simultaneous treatment with PMSF does not exhibit any signal decrease. GA and 643 PMSF treatment shows a similar amyloid signal after 2 hours, while ABA and endopeptidase treatment 644decreases it. DLS analysis of the SSPB with the proteases with or without PMSF are represented in (m-645 n). The treatment with proteases decreases the size of the SSPB, while presence of PMSF prevents the 646 decrease in size. (Gamma value for each image is 1.3-1.6, Graphs are represented as mean with error 647 bars as standard error of mean, Scale bars for microscopy corresponds to 50 µM)

Now, to confirm the role of the signalling molecules on the amyloid content of the protoplasts, these were treated with GA, ABA and an equimolar mixture of GA and ABA. ThT staining was performed and imaged immediately and after 2 hours. Wheat aleurone (**Figure 5 a1-a4)** and mungbean cotyledon **(Figure 5 c1-c4)** protoplasts treated with incubation buffer, GA, ABA, GA and ABA respectively at 0 hour show ThT signatures in the SSPB. The wheat (**Figure 5 b1-b4)** and mungbean (**Figure 5 d1-d4)** protoplasts after treatment for 2 hours, were again visualized for changes in ThT signature. The GA-treated protoplasts show a decrease in ThT fluorescence **(Figure 5 b2, d2, i and j)** for both wheat and mungbean, suggesting GA-regulated decrease in amyloid content. The ABA treated protoplasts show a similar intensity **(Figure 5 b3, d3, i and j)** as compared to control, whereas treatment with the mixture decreases the intensity in wheat but not in mungbean, reflecting on the possible differences in the amyloid structures of these two seeds **(Figure 5 b4, d4, i and j)**.

After establishing the role of the plant hormones on amyloid degradation, the effect of proteases on these structures was investigated. At first, the seed endopeptidase fraction was isolated and purified. For this purpose, the mungbean and wheat seed **(Figure S8 a1-a2)** total protein isolates were analysed by size-exclusion chromatography and the fractions collected were checked for protein estimation. The protein-containing fractions were checked for enzymatic activity on casein as a substrate. Purified papain, was used as the control since papain-like proteases are upregulated in the seeds during germination.(Liu *et al*., 2018) The representative enzymatic activity of the fractions from the 48h imbibed seeds and papain are shown in **Figure S8 a3-a4**. The isolated fractions were then checked for their time-dependent activity on casein and at each time-point, the amount of amino acids released due to digestion was assayed by ninhydrin, using casein as a substrate. The activity of the endopeptidases isolated from mungbean and wheat **(Figure S8 a5-a6)** after 48 hours of imbibition was found to be optimum.

The isolated protoplasts were next treated with papain (control) and endopeptidase fraction with or without PMSF. The wheat **(Figure 5 e1, f1)** and mungbean **(Figure 5 g1, h1)** protoplasts incubated with either papain, or endopeptidase fraction, lose their amyloid signature significantly over a period of 2 hours (**Figure 5 e2, f2, g2 and h2)**. When treated with both papain and PMSF, **(Figure 5 e3, f3, g3 and h3)** or endopeptidase and PMSF **(Figure 5 e4, f4, g4 and h4)**, the signature was retained, showing that PMSF inhibits the proteases. When the protoplasts were incubated with GA and PMSF, interestingly the signature is still retained, while GA alone leads to loss of amyloid signatures. This confirms that GA-induced upregulation of endoproteases is countered by the presence of PMSF **(Figure 5 e5, f5, g5 and h5)**. Lastly, when the protoplasts were incubated with ABA and endopeptidase, **(Figure 5 e6, f6, g6 and h6)** the signatures lose, indicating that although ABA might be counteracting the effect of endogenous GA inside the protoplast, it cannot counter the effect of exogenous endopeptidase. The quantification is further confirmed by ImageJ **(Figure 5 k-l)**. A similar pattern is observed in the CR-stained protoplasts **(Figure S9)**. DLS analysis of the SSPB incubated with papain, and the endopeptidase fraction further exhibits the degradation of the SSPB, as evident from the decrease in the hydrodynamic diameter, while the presence of PMSF leads to maintenance of SSPB size **(Figure 5 m-n)**.

Summarizing these results, the *in-situ* and *ex-vivo* results indicate that a systematic degradation of the amyloid-containing SSPB is controlled by hormones and their downstream effectors, for facilitating germination.

## Discussion

The proteinaceous plant SSPB is known for its role in germination. However, the complex internal arrangement of proteins is not elucidated till now. In morphological aspects, SSPB are similar to other organisms’ amyloid-containing protein bodies. Also, previously, some plant proteins were shown to form amyloid fibrils *in-vitro* under non-native conditions. Based on these, we hypothesized that the monocot and dicot SSPB might contain similar structures *in-vivo* for modulating seed physiological functions. Importantly, experimental validation using multiple amyloid-specific dyes is imperative to assign amyloid or amyloid-like properties to proteins and to remove the prevailing perplexity of the presence of proteinaceous amyloids and glucan structures.(Matiiv *et al*., 2020) To establish this, we employed a multispectral strategy and have overcome these issues using several amyloid-specific probes. Fascinatingly, whereas ThT and Proteostat^®^ bound to the entire SSPB-containing regions of aleurone and cotyledon cells, the birefringence signals of amyloid due to CR staining appeared in only some areas. The amyloidogenic proteins were confirmed by LMD-MS/MS and the amyloidogenicity is allotted primarily to the globulins.

To further establish the hypothesis and to gain an insight into the amyloidogenic properties of these structures, we isolated the SSPB from the aleurone and cotyledon cells and utilized these for physicochemical studies including DLS, SEM, TEM, HRTEM, FTIR, SDS-PAGE and XRD. The SSPB and the dialyzed globulin fibrils exhibit ThT and CR signatures, prominent amyloid signatures in IR spectra and fibrillar structures in HRTEM. These also exhibit detergent resistance and amyloid-specific reflections in XRD. On the other hand, the isolated protein fractions from the SSPB in their soluble state (globulin and albumin), are devoid of such signatures, and suggests that they assemble in the SSPB and exhibit amyloid composite structure.

Next, to assess the functional roles of the amyloid structures in seed, their signatures and content was followed during the germination and seedling growth by *in-situ* techniques. The results suggest that the ThT-stained amyloid-like structures degrade faster compared to the CR-stained amyloids, due to the differential stability of these two assemblies. The SSPB isolated from germinated seeds further show a decrease in the propensity of amyloid signature by IR and establishes the degradation of the amyloids in germination. The amyloidogenic protein content of the CR positive amyloid areas after 72 hours in mungbean and wheat further shows that some storage proteins degrade slower compared to others and might be present in more stable structural arrangement.

Since seed germination is dependent on plant hormones and proteases; we next carried out functional assay of the amyloids by employing exogenous hormones, proteases and their inhibitors. These results were further validated using protoplasts, which serve as the active metabolic state of the seed cotyledon and aleurone cells. The protoplasts were utilized for investigating the molecular players responsible for amyloid degradation during germination. The results indicate that on seed imbibition, GA would lead to an enhanced expression of proteases which can subsequently degrade the amyloid structures present in the SSPB. The effect can be countered with the help of either ABA, a known antagonistic molecule of GA, or by using protease inhibitors such as PMSF. The inhibition of amyloid degradation was found to reduce the germination index, further confirming that amyloid degradation plays a crucial role for controlling seed germination in the wheat and mungbean seeds.

Since amyloid formation follows pathways of nucleation and elongation, the same phenomenon may explain the biogenesis of the SSPB, as aggregated structures by self-seeding and cross-seeding of other protein components. The presence of these structures might facilitate protection of seeds from various environmental stresses. The current study would open up new research questions to study the SSPB biogenesis and might illuminate the enigmatic issue of the role of functional amyloids in evolution across species.

## Supplemental material

### Supplementary Section 1

- **Cellulase treatment and Congo red/ThT staining**,
- **Isolation of protein fractions and physicochemical characterization of SSPB**,
- **Amyloidogenic properties of SSPB and the isolated proteins**,
- **Isolation of endoprotease from the germinated seeds and sustained release**
- **Treatment of the seeds and protoplasts with exogenous molecules and their inhibitors**,
- **Processing of LMD samples and protein fractions for MS/MS**

1. **Movie S1 Congo red birefringence in aleurone cells of barley**
2. **Movie S2 Congo red birefringence in cotyledon cells of chickpea**
3. **Fig. S1 Low magnification images of ThT-stained seeds and control images for amyloid-specific staining**
4. **Fig. S2 Demarcation of glucan-rich regions and amyloidogenic content analysis of major storage proteins of seeds**
5. **Fig. S3 Physicochemical characterization of the SSPB and the reconstituted fraction**
6. **Fig. S4 The amyloidogenic properties of the isolated protein fractions**
7. **Table. S1 MS/MS analysis and top-scoring proteins in the SSPB protein fractions**
8. **Fig. S5 ThT and CR fluorescence of germinated seeds to detect amyloids vs amyloid-like aggregates, biophysical characterization of SSPB from germinated seeds and sustained release**
9. **Fig. S6 Quantitation of the proteinaceous signal in the germinated seed sections**
10. **Fig. S7 The presence of amyloid-like, amyloids and protein content at different time-points of germination**.
11. **Fig. S8 Seed endopeptidase characterization and *in-vitro* biological activity**
12. **Fig. S9 Congo red staining of protoplasts treated with exogenous molecules and protoplast control for isolation and staining**
13. **Table S2: List of the number of peptide sequences identified for each protein identified for mungbean and wheat**

## Acknowledgements

We are thankful to Prof. Pradip Sinha and Prof. Amitabha Bandyopadhyay for letting us use their confocal imaging facility and ultramicrotome facility. We also thank Mr. Upendra Singh Yadav and Mr. Saurabh Singh Parihar for helping with these facilities. We are thankful to Dr. Geetashree Mukherjee and Mr. Suman Sarkar for the laser capture microdissection facility and Dr. Saravanan Matheshwaran for size exclusion chromatography. We are thankful to Advanced Imaging Centre IIT Kanpur and Mr. Jai Singh for TEM and HRTEM analysis. We are thankful to Dr. Shraddha Singh and BSBE department for SEM analysis and to IITK-ACMS X-ray diffraction facility for powder diffraction data. N.S and T.Z. acknowledge University Grants Commission for Senior Research Fellowship.

## Data availability

The data that support the findings of this study are available from the corresponding author upon reasonable request.

## Author contributions

The original concept was conceived by A.K.T. two decades ago. A.K.T supervised the experiments; A.K.T. and N.S designed the experiments. N.S. carried out most of the experiments conducted in the current manuscript. A.K.T and N.S. analyzed the data; A.Y.G performed the initial sectioning and staining experiments; N.S. and T.Z. performed the sectioning, staining and protoplast isolation experiments. B.R. independently validated staining data and performed SDS-PAGE. A.B., S.B. and A.C. helped in proteomics data acquisition and analysis. A.K.T and N.S. wrote the manuscript.

