## Supplemental Data for "Protein reservoirs of seeds are composites of amyloid and amyloid-like structures facilitating sustained release during germination and seedling growth"

*New Phytologist* Supporting Information

Article acceptance date: Click here to enter a date.

The following Supporting Information is available for this article:

**Supplementary Section 1:**

- **Cellulase treatment and Congo red/ThT staining,**
- **Isolation of protein fractions and physicochemical characterization of SSPB,**
- **Amyloidogenic properties of SSPB and the isolated proteins,**
- **Isolation of endoprotease from the germinated seeds and sustained release,**
- **Treatment of the seeds and protoplasts with exogenous molecules and their inhibitors**
- **Processing of LMD samples and protein fractions for MS/MS**

**1. Movie S1 Congo red birefringence in aleurone cells of barley**

**2. Movie S2 Congo red birefringence in cotyledon cells of chickpea**

**3. Fig. S1 Low magnification images of ThT-stained seeds and control images for amyloid-**

**specific staining**

**4. Fig. S2 Demarcation of glucan-rich regions and amyloidogenic content analysis of major storage proteins of seeds**

**5. Fig. S3 Physicochemical characterization of the SSPB and the reconstituted fraction**

**6. Fig. S4 The amyloidogenic properties of the isolated protein fractions**

**7. Table. S1 MS/MS analysis and top-scoring proteins in the SSPB protein fractions**

**8. Fig. S5 ThT and CR fluorescence of germinated seeds to detect amyloids vs amyloid-like aggregates, biophysical characterization of SSPB from germinated seeds and sustained release**

**9. Fig. S6 Quantitation of the proteinaceous signal in the germinated seed sections**

**10. Fig. S7 The presence of amyloid-like, amyloids and protein content at different time-points of germination.**

**11. Fig. S8 Seed endopeptidase characterization and *in-vitro* biological activity**

**12. Fig. S9 Congo red staining of protoplasts treated with exogenous molecules and protoplast control for isolation and staining**

**13. Table S2: List of the number of peptide sequences identified for each protein identified for mungbean and wheat**

**Supplementary Section 1:**

**Materials:** The seeds (cereals-wheat, barley, rice, maize; pulses – chickpea, mungbean, soybean, fieldpea) were acquired from ICAR-Indian institute of pulses research, and Chandra Shekhar Azad University of Agriculture and Technology, Kanpur, India and stained with amyloid-specific probes. Although all the seeds chosen showed similar results, in the current paper, the best representative seeds (wheat, barley, chickpea and mungbean) are presented.

**a. Cellulase treatment and Congo red/ThT staining:** To ensure that the birefringence observed by Congo red probe is due to the amyloid content and not due to the glucan structures, the seed sections were incubated in 4.5% cellulase solution for 10-120 minutes in mungbean and 10 minutes to 72 hours in wheat. The sections were then simultaneously stained with Congo red and Calcofluor white. Sections without cellulase treatment were used as control. Separate sections were used for simultaneous staining of ThT and Calcofluor white to ensure that the cell wall regions and the specified amyloid/amyloid-like regions are distinct and there are no overlaps.

**b. Isolation of protein fractions from SSPB and physicochemical characterization of SSPB:**

For water-soluble albumin and salt-soluble globulin fraction extraction, previously established protocols were followed, using the isolated SSPB as the source of the protein fractions. For albumin, SSPB (100,000 per ml) were lysed with a double amount of lysis buffer and stirred for 2 hours in water containing 10 mM calcium chloride, 10 mM magnesium chloride and 0.5% of Triton-X. The insoluble proteins were pelleted by centrifugation at 30000g at 4˚C for 1 hour. The supernatant was collected as the water-soluble albumin fraction. For isolating globulin, the pellet was resuspended in Tris-HCl (10 mM) buffer at pH 7.5 containing 10% w/v NaCl, 10 mM EDTA, and stirred for 2 hours at 4˚C. Centrifugation was performed at 30000 g at 4˚C for 1 hour to separate globulins, which are salt-soluble and remain in the supernatant.(Czubiński *et al.*, 2016) The protein fractions were purified using 40% ammonium sulfate precipitation. Nanodrop was used to check contamination of non-protein molecules, including nucleic acids (an absorbance peak at 260 nm or the ratio of A260/A280 more than 1 signifies DNA/RNA contamination in the protein sample). For reconstitution studies, 12 kDa and 3.5 kDa dialysis membranes were used for desalting the globulin and albumin fraction, respectively. Dialysis was repeated thrice for each type of protein to ensure maximum salt removal.

For **scanning electron microscopy (SEM)** analysis of SSPB, the isolated SSPB (10000 per ml) in 1X PBS were spotted on coverslips and fixed with 0.3% glutaraldehyde for 30 minutes. After fixation, the coverslips were treated with increasing gradients of ethanol

(50-100%) to dehydrate the samples. The tissue and SSPB samples were gold-coated and analysed by Carl Zeiss EVO18 SEM. For **transmission electron microscopy (TEM) and high resolution-TEM (HRTEM)** analysis, the isolated intact SSPB (10000/ml) and 0.5% Triton-X treated SSPB (for disrupting SSPB) in 1X PBS (after similar treatment as that of SEM) were spotted on carbon-coated copper grids and stained for 20 seconds with 1% uranyl acetate and analysed using FEI-Tecnai Twin 120 KV TEM and FEI-Titan G2 60-300 KV HRTEM. The method utilized for SEM and TEM analysis were adopted and modified from previously established protocols.(Antonets *et al.*, 2020) For **bright field/fluorescence** microscopy analysis, the SSPB (10000/ml) were treated with 20 μM ThT or 1 mg/ml CR solution in water for 15 minutes.

For confirming the amyloid signatures in the SSPB, these were isolated at different time-points of germination and were checked for **ThT fluorescence** at 450/489 nm. The number of SSPB was normalized at each time-point to negate the effect of decrease in SSPB number with germination (5000 SSPB/ml for each time-point). For **FTIR** analysis, a Bio ATR-FTIR Bruker Tensor IR-27 instrument was used. 20 μL of each sample (0.5 mg/ml) was added to liquid nitrogen cooled ZnSe crystal and scanned from 800-4000 cm$^{-1}$. The secondary derivatives were calculated to find the peaks and Gaussian fit was performed using Origin Pro 9.1 software. The amide I bands (1600-1700 cm$^{-1}$) for each spectra was then analysed for secondary structure content. The absorbance (arbitrary units) in each spectrum was simply used to gauge the relative predominance of a particular secondary structure and was not used for further analysis. In all the FTIR spectra represented, the overall fitted curve is shown in black and each individual peak is assigned a different colour (red, green, blue and cyan from left to right). The peak wavenumbers are presented adjacent to each peak. **DLS analysis** was performed on Malvern Zetasizer ZS90 using a scatter angle of 90°. For each sample, 0.5 mg/ml protein solution was used.

**Amyloidogenic properties of SSPB and the isolated proteins:** The isolated globulin proteins were dialysed and during different time-points of dialysis, aliquots were incubated with 20 μm ThT solution and were analysed for their fibrillation kinetics. The dialysed protein samples were analysed by TEM to confirm their fibrillar morphology.

To decipher detergent resistance of SSPB fibrils and the *in-vitro* formed fibrils of dialyzed globulin, the SSPB were disrupted to remove the lipid membrane and the fibrils were boiled with SDS for 10-120 minutes prior to performing SDS-PAGE to show comparative migration (12% polyacrylamide gel)(Pan & Zhong, 2015; Song *et al.*, 2016). Similarly SDS-PAGE was performed the dialysed globulin fibrils and albumin protein fractions. To further confirm the amyloidogenic nature of the *in-vitro* formed fibrils of globulin and SSPB, the samples were lyophilized and analysed by powder X-ray diffraction (Pananalytical X'Pert Powder Diffraction, X-ray wavelength – 0.154 nm, 2θ range 5-60˚) to check presence of cross-β-sheet structures, a characteristic of amyloids.

**c. Isolation of endoprotease from the germinated seeds and sustained release:** For this purpose, the total protein fraction was isolated from the seeds (0, 24 and 48 hours imbibed) according to previously established protocols and purified with ammonium sulphate precipitation (4ºC). (Görg *et al.*, 2006) The protein sample (1 mg/ml, injection volume 500 µl) in 10 mM Tris-HCl buffer, pH 8, was analysed by size-exclusion chromatography using a Superdex 200 column with 0.5 ml/minute flow-through rate (4ºC). The column was equilibrated with 10 mM Tris-HCl buffer, pH 8. For calibration a pre-made standard curve was used (1.5-600 kDa). The fractions were collected and each fraction was analysed for protein concentration using bicinchoninic acid (BCA) assay, using bovine serum albumin (BSA) as a standard. The fractions with protein were checked for their enzymatic activity. For this, the protein-containing fractions were incubated with casein (2%), a standard substrate at 25˚C, pH 6.5. At fixed time-points, the proteins were precipitated with 1% tri-chloro acetic acid and centrifuged at 6000 rpm for 15 minutes. The supernatant was checked for amino acid content using ninhydrin assay, by measuring absorbance at 570 nm using Perkin Elmer micro-plate reader.(Silveira *et al.*, 2017)

**d. Treatment of the seeds and protoplasts with exogenous molecules and their inhibitors:** For sustained release assay, each substrate (albumin, globulin, their dialyzed aggregates, SSPB and SSPB without membrane) and the endopeptidase fraction (0.5 mg/ml) were incubated together in the upper chamber of a 200-nm trans-well filter, as commonly employed for checking sustained degradation of drugs and other biomolecules.(Rohrschneider *et al.*, 2015) The filters were placed in the wells of a 24-well plate and dipped in 10 mM Tris-HCl, pH 8. At each time-interval, the samples were withdrawn from the lower chamber, precipitated with 1% tri-chloro acetic acid and analysed by ninhydrin assay for quantifying the amino acids released.

For functional relevance of the amyloids in seed germination, wheat and mungbean seeds were imbibed in water, gibberellin (GA), GA and phenyl methyl sulfonyl fluoride (PMSF), abscisic acid (ABA), ABA and PMSF, and PMSF alone for 72 hours. All the treatments were added at a final concentration of 50 μM. The seeds were then fixed, processed, and CR-ThT staining was performed to check amyloid retention. The monocot and dicot protoplasts were treated with incubation buffer, GA, ABA, GA and ABA, papain, seed endopeptidase, papain and PMSF, endopeptidase and PMSF, GA and PMSF, ABA and endopeptidase at the same final concentration of 50 μM for 2 hours. The protoplasts were stained with ThT or CR and visualized to check amyloid signals. Each experiment with protoplasts was repeated at least thrice (n=3) with freshly prepared protoplasts.

**e. Processing of LMD samples and protein fractions for MS/MS**

The collected LMD samples from CR, ThT and acid-fucshin stained seed sections, and isolated SSPB were centrifuged at 6000 rpm for 10 minutes. To each tube, 40 µl of lysis buffer (10 mM Tris, 1 mM EDTA and 0.5% Triton-X 100) was added to aid protein extraction. The samples were sonicated in a bath sonicator for 20 minutes followed by denaturation by incubating at 95ºC for 1 hour. The albumin and globulin fractions, isolated from SSPB, were denatured at 95ºC for 1 hour. The protein concentration in each sample was measured using BCA assay and 30 µg of protein of each sample was isolated for digestion. To every sample, 0.5 µg of trypsin gold was added and digestion was continued for 18 hours at 37ºC. The peptides generated, were reduced by using 5 mM dithiothreitol for 30 minutes at 37ºC. This was followed by alkylation with 15 mM iodoacetamide in dark for 45 minutes. The samples were centrifuged at 20000 g for 20 minutes. The clear supernatant was collected and dried in vacuum desiccator at 30ºC for 1 hour. The samples were resuspended in 0.1% trifluoroacetic acid (25 µl) and desalinated using C18 ZipTip columns. The samples were finally resuspended in buffer and injected in the autosampler of the Ekspert Nano LC 425 connected to AB Sciex Triple TOF 6600 mass spectrometer for LC-MS/MS.

The wheat spectra obtained were analysed using SciEX software against the available database. For mungbean, due to unavailability of the premade database, a new library was prepared from UniProt. PeptideShaker and ProteinPilot were used to analyse the mungbean spectra. For each dataset, although we obtained multiple proteins with high confidence score, we have represented the top 4 proteins for each sample.

189

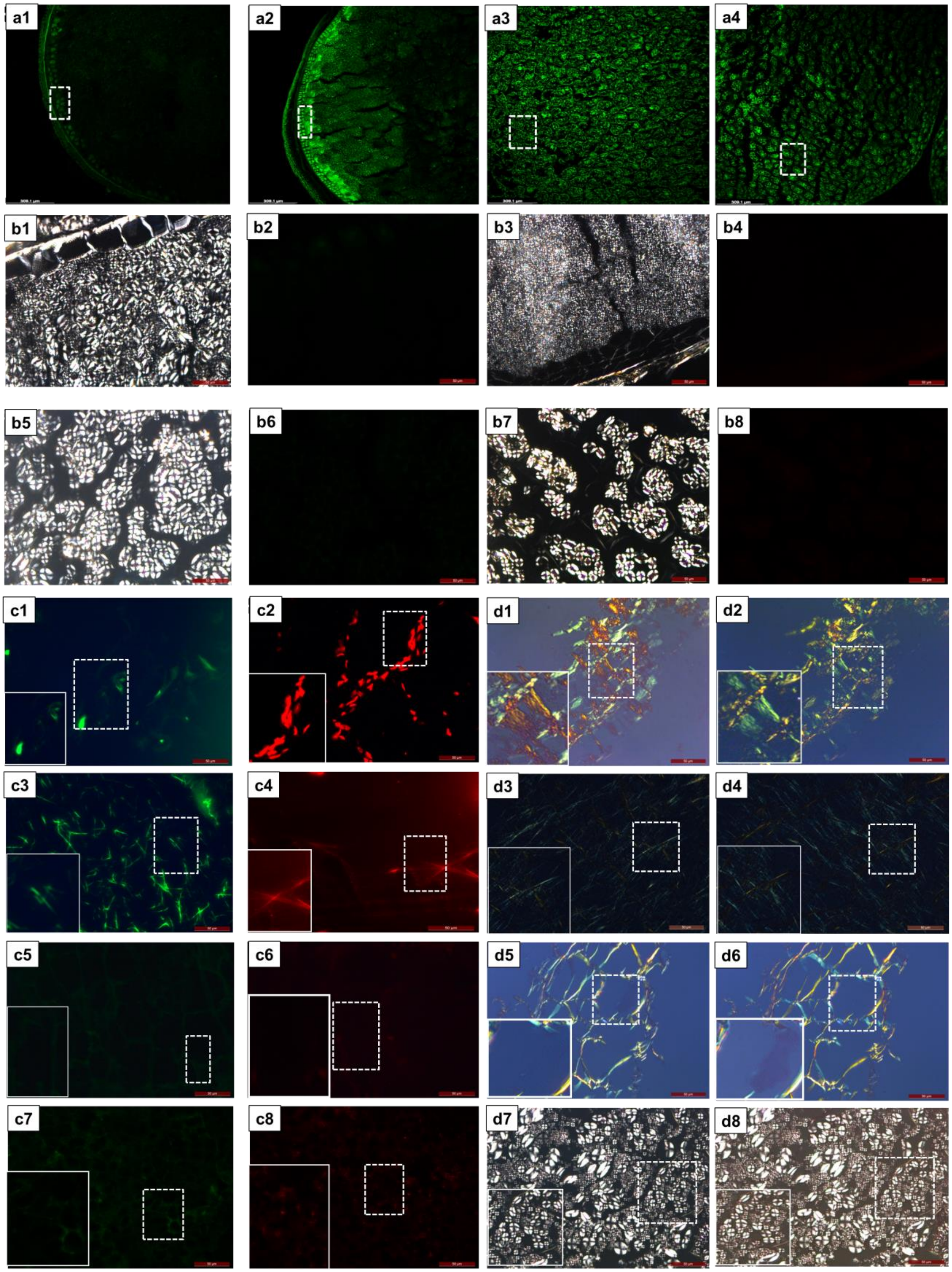
a1
a2
a3
a4
b1
b2
b3
b4
b5
b6
b7
b8
c1
c2
d1
d2
c3
c4
d3
d4
c5
c6
d5
d6
c7
c8
d7
d8

**Fig. S1 Low magnification images of ThT-stained seeds and control images for amyloid specific staining**

The 10X images of ThT stained seed sections of wheat, barley, chickpea and mungbean are represented (a1-a4). The white dashed boxes represent the intense ThT-positive areas. Unstained sections of wheat (polarizer – b1, GFP filter – b2), barley (polarizer – b3, GFP filter – b4), chickpea (polarizer – b5, GFP filter – b6) and mungbean (polarizer – b7, GFP filter – b8) showing neither birefringence nor fluorescence. As positive controls of ThT and Proteostat®, human abdominal fat biopsy tissue of a positive amyloidosis patient (c1, c2) and amyloidic fibrils of Sup35-N-terminal domain peptide (GNNQQNY) is represented (c3, c4), showing intense green or red fluorescence due to positive staining. As negative controls of the two probes, sections of potato tuber as a storage organ (c5, c6) and the wheat endosperm is represented (c7, c8) where, negligible fluorescence is observed. Gamma values for each ThT and Proteostat® image ranges from 1.8-2.0 to bring uniformity, changes in brightness/contrast have been applied to the whole image. As positive controls of CR staining, human abdominal fat biopsy tissue of a positive amyloidosis patient (d1, d2) and amyloidic fibrils of Sup35-N-terminal domain peptide (GNNQQNY) is represented (d3, d4), exhibiting green-to-red birefringence. As negative controls of the probe, sections of potato tuber as a storage organ (d5, d6) and the wheat endosperm is represented (d7, d8), where negligible birefringence is observed. The white dashed boxes show specific areas and solid lined boxes show the magnified portions of these areas. Gamma values are at 1.0 for facilitating clear birefringence visualization. Scale bar – 50 μm.

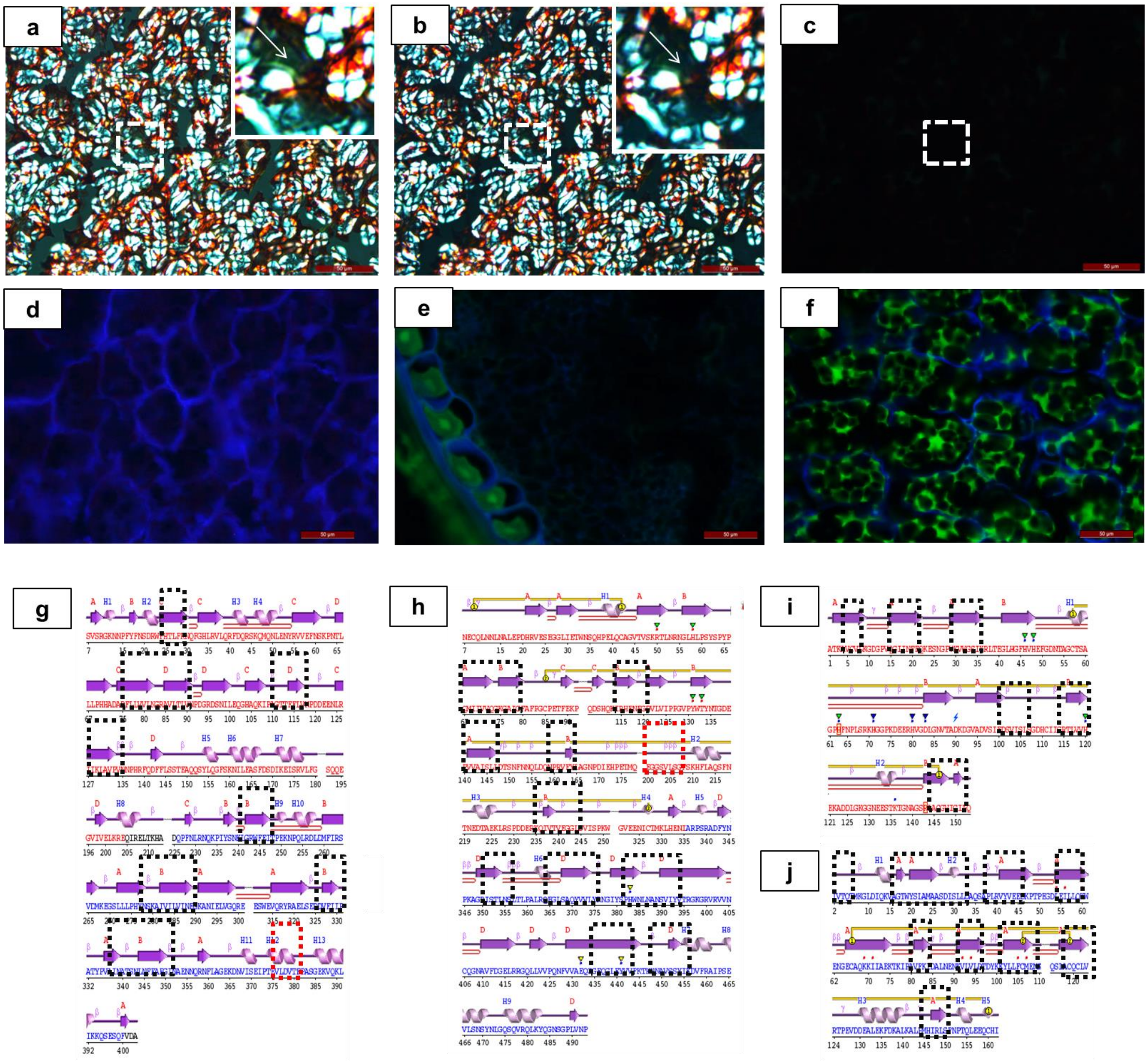


| k Protein | Uniprot ID | Number of hotspots | Total hotspot area % |
|---|---|---|---|
| Mungbean 8S Globulin | Q198W4 | 12 | 45.4 |
| Mungbean 2S Albumin | A0A1S3VUC6 | 2 | 17.7 |
| Chickpea Vicilin-like globulin | A0A1S2XQR4 | 11 | 40.4 |
| Chickpea 2S Albumin-like | A0A1S2XDF0 | 2 | 18.4 |
| Soybean Glycinin G1 Globulin | P04776 | 14 | 43.3 |
| Soybean Albumin-1 | Q39837 | 5 | 22.7 |
| Wheat Globulin-3 | B7U6L4 | 14 | 35.7 |
| Maize 7S globulin | A0A3L6E9P2 | 21 | 52.7 |
| Maize Albumin | ALB3 | 8 | 22.9 |

**Fig. S2 Demarcation of glucan-rich regions and amyloidogenic content analysis of major storage**

**proteins of seeds**

The binding of Congo red to amyloids is confirmed after cellulase treatment. Even after cell wall lysis, the presence of Congo red birefringence indicates the presence of amyloids in mungbean (a-b). A complete lack of Calcofluor white signal in (c) confirms cell wall digestion. Comparatively, the strong blue fluorescence in (d) in another section with no cellulase treatment, shows presence of cell walls. Simultaneous staining with both ThT and Calcoflour white suggests that ThT binds to only amyloidic regions. Moreover, presence of glucan structures are not observed in the ThT-positive regions in these sections, confirming non-overlapping spatial distribution of amyloids and glucans.(e-f) Secondary structure content of mungbean 8S and soybean 11S globulin respectively (g-h), the same is represented for amyloidogenic β-barrel containing human superoxide dismutase and bovine lactoglobulin respectively (i-j). The black and red dashed boxes indicate potential aggregation-prone regions as predicted by AggreScan web server. Black boxes represent the aggregating regions present in the β-strands while red boxes represent the same in other secondary structures. The dicot and monocot globulins and albumins are next enlisted with the number of hotspots and total hotspot area (k). The analysis suggests that globulins are more prone to aggregation as compared to albumins.

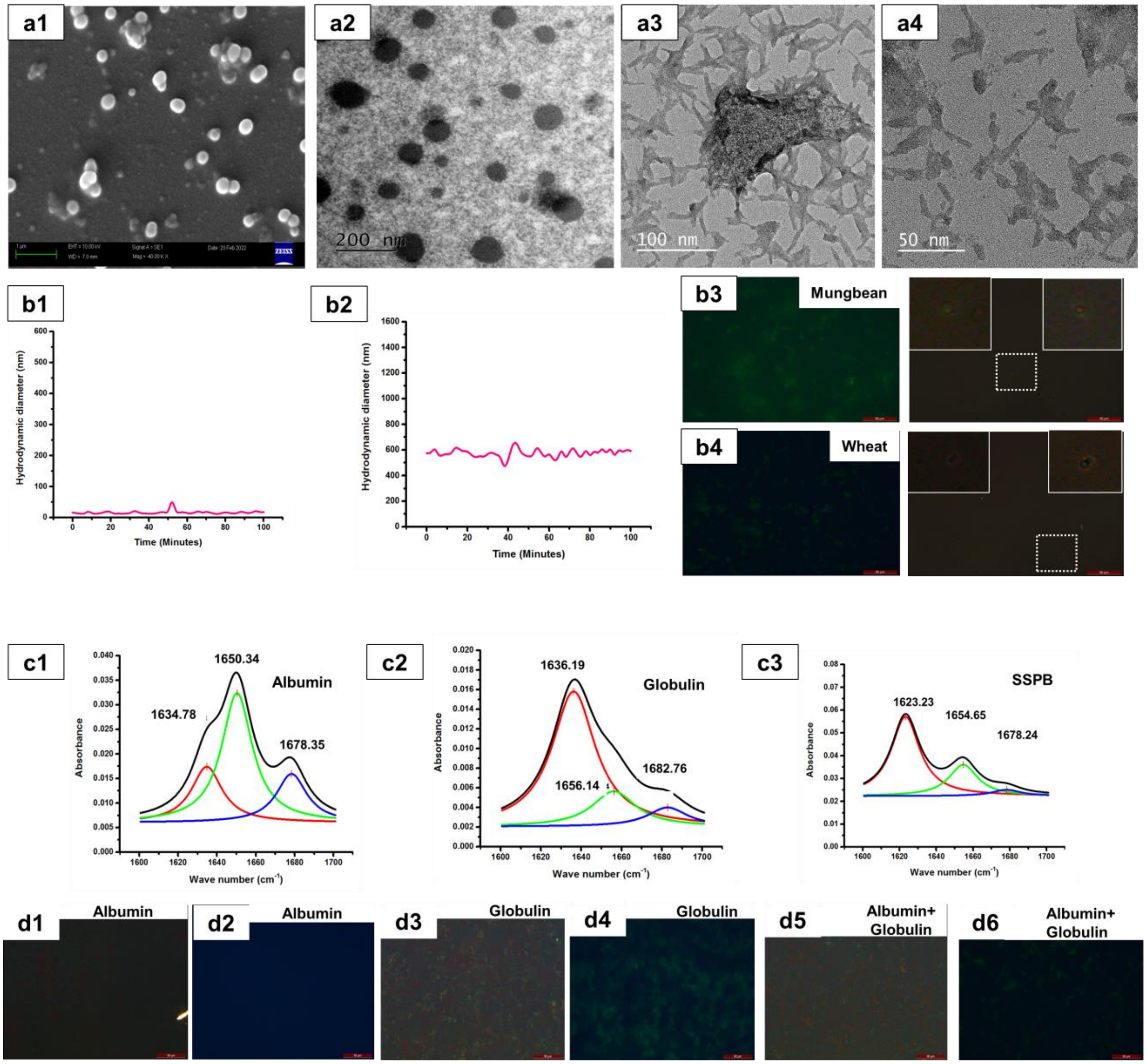


**Fig. S3 Physicochemical characterization of the SSPB and the reconstituted fraction**

SEM (a1) and TEM (a2) analysis of isolated SSPB of wheat reveals mostly spherical structures. On disrupting the SSPB, fibrils are observed in HRTEM (a3-a4). DLS analysis of the SSPB of mungbean (b2) and wheat (data not shown) confirm similar hydrodynamic diameter as observed in SEM, whereas the soluble globulin fraction (b1) shows a much smaller diameter. ThT and CR stained SSPB of mungbean (b3) and wheat (b4) show signal with both the probes, confirming their amyloidic nature. The left images represent ThT, whereas the right images represent CR staining. FTIR analysis of the secondary structure content of water-soluble albumin, salt-soluble globulin fraction and the SSPB of wheat (c1-c3) reveal that albumin consists of helical, sheet and turn structures while globulins have predominant β-sheet signatures as evident by the major peak at 1636 $cm^{-1}$. The SSPB on the other hand, shows evidence of amyloids by the shift towards 1623 $cm^{-1}$. Dialyzed albumin fraction of mungbean does not show any CR (d1) or ThT (d2), but dialyzed globulin exhibits intense CR (d3) and ThT (d4) signals. Dialyzed globulin and albumin when mixed together (4 hours), show similar signatures (d5-d6). (Scale bar of SEM -1 µm, TEM – 200 nm, HRTEM – 100 and 50 nm)

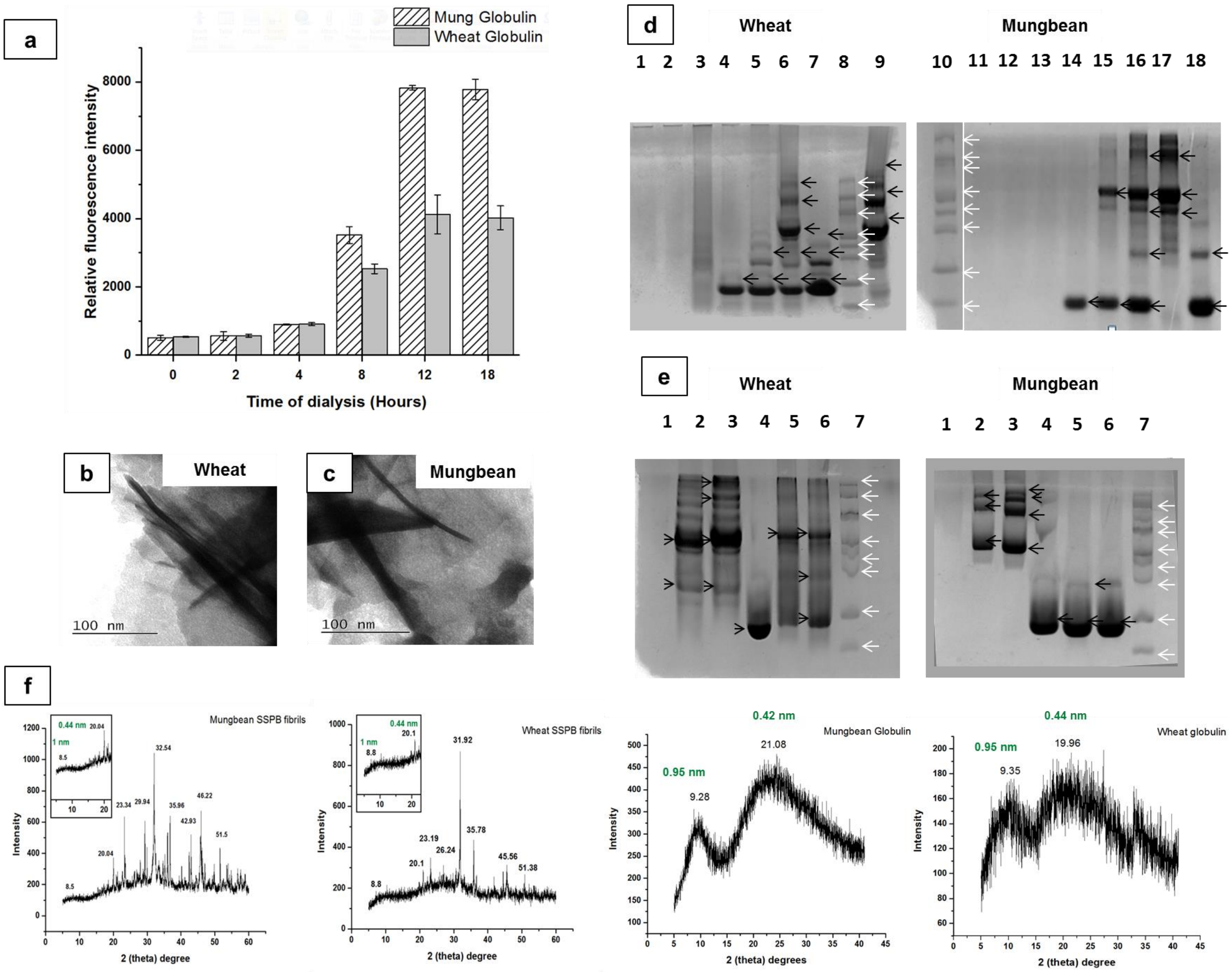


**Fig. S4 The amyloidogenic properties of SSPB and isolated protein fractions.** The fibrillation of the globulin proteins during different time points of dialysis using ThT (a). Fibrillar amyloid structures of dialyzed globulin fractions of wheat and mungbean (b-c). Detergent resistance of SSPB [1,11- Undisrupted unboiled SSPB, 2,12- Disrupted unboiled SSPB, 3,13- Undisrupted boiled SSPB for 10 minutes, 4,14- disrupted boiled SSPB for 10 minutes, 5,15 – disrupted boiled SSPB for 60 minutes,6,16 – disrupted boiled SSPB for 120 minutes, 7,18- albumin protein, 9,17 – globulin proteins and 8,10- Protein Ladder] of wheat and mungbean (d), Detergent resistance of dialysed globulin and albumin [1 – Globulin boiled for 10 minutes, 2- Globulin boiled for 60 minutes, 3- Globulin boiled for 120 minutes, 4- albumin boiled for 10 minutes, 5-albumin boiled for 60 minutes, 6-albumin boiled for 120 minutes].(e) X-ray powder diffraction of SSPB fibrils and globulin fibrils of wheat and mungbean (f) Ladder

**Table. S1 MS/MS analysis and top-scoring proteins in the SSPB protein fractions.** The proteins of the water-soluble and salt-soluble fractions of mungbean and wheat are represented.

| Mungbean salt-soluble fraction | Mungbean water-soluble fraction | Wheat salt-soluble fraction | Wheat water-soluble fraction |
|---|---|---|---|
| 8S Globulin alpha isoform | Mungbean seed albumin | Globulin 3A | Ubiquitin |
| 8S Globulin beta isoform | Class I heat-shock protein | High molecular weight glutenin | Genome assembly protein |
| Basic 7S Globulin-2-like | Actin | Globulin 1 | Glyceraldehyde 3 phosphate dehydrogenase |
| Vicilin-like seed storage protein | Alcohol dehydrogenase | Globulin 3B | Actin |

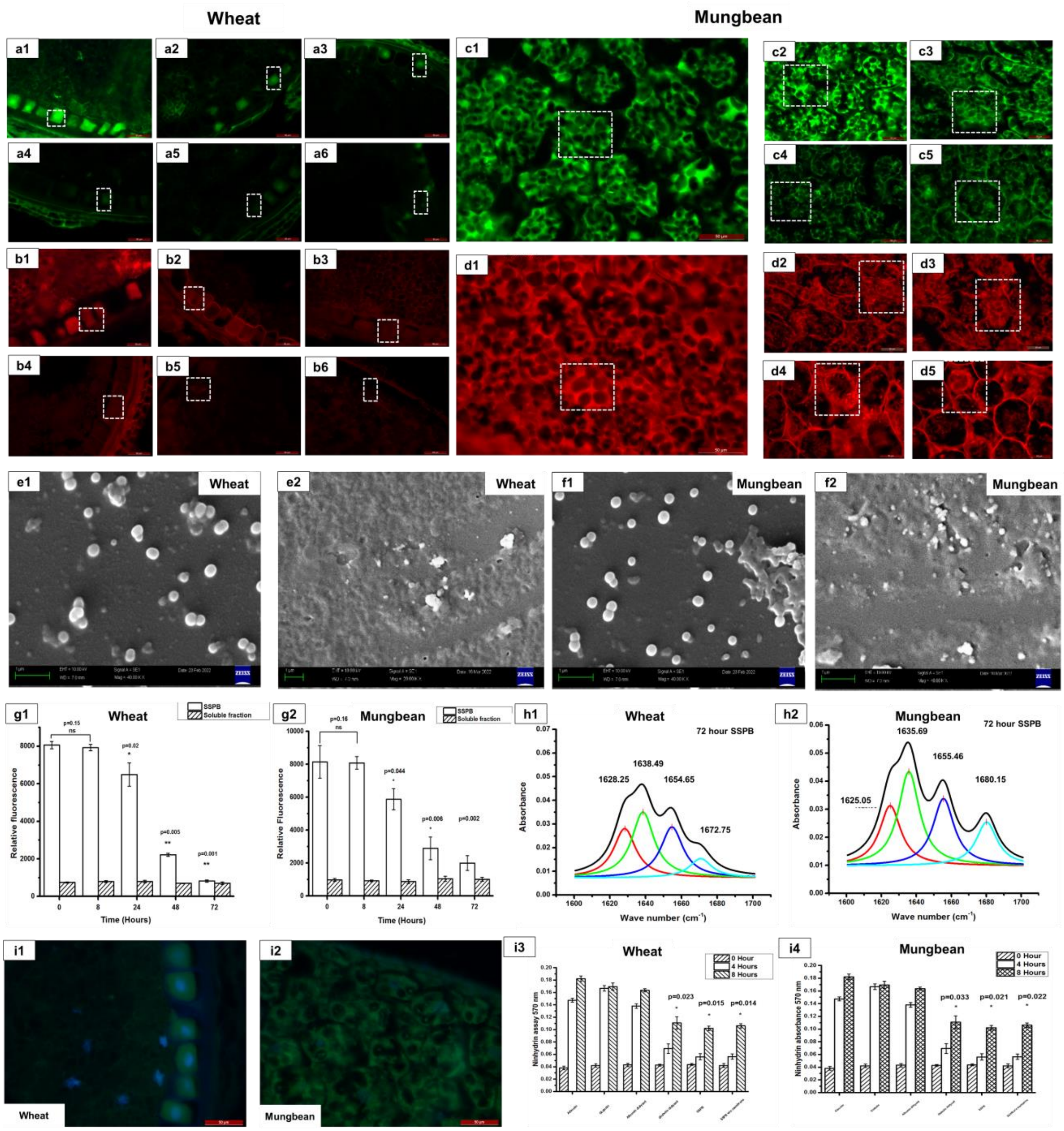


**Fig. S5 ThT and CR fluorescence of germinated seeds to detect amyloids vs amyloid-like aggregates, biophysical characterization of SSPB from germinated seeds and sustained release**

ThT staining of wheat seeds are represented at 0 (a1), 8 (a2), 24 (a3), 48 (a4), 72 (a5) and 96 (a6) hours. CR staining of wheat seeds are represented at 0 (b1), 8 (b2), 24 (b3), 48 (b4), 72 (b5) and 96 (b6) hours. ThT staining of mungbean seeds are represented as 0 (c1), 8 (c2), 24 (c3), 48 (c4) and 72 (c5) hours, and CR staining of mungbean seeds are represented as 0 (d1), 8 (d2), 24 (d3), 48 (d4) and 72 (d5) hours. ThT signal in wheat start decreasing since 8 hours and in mungbean from 24 hours. The CR fluorescence however decreases after 72 hours in wheat and does not decrease significantly till 72 hours in mungbean. The white dashed boxes represent representative areas. (Gamma value for each image is 1.3-1.4) (Scale bar – All scale bars correspond to 50 μm). SEM analysis of the SSPB isolated from 0 hour and 72 hour germinated wheat (e1-e2) and mungbean (f1-f2) seed, show a decrease in size and number. ThT fluorescence of wheat (g1) and mungbean (g2) SSPB and soluble globulin fraction with germination, indicate the decrease in amyloidic content of the SSPB while the soluble fraction's fluorescence is maintained at a constant baseline level. FTIR signatures of SSPB at 72 hours of germination for wheat (h1) and mungbean (h2) show that with time, the decrease in the inter-sheet interactions in the SSPB is evident along with an increase in helices and other disordered structures, suggesting the disappearance of amyloidic character of the SSPB with germination. ThT and DAPI stained representative images of wheat and mungbean (i1-i2). Sustained release of amino acids of SSPB and amyloidic aggregates, compared to the soluble proteins using ninhydrin assay (i3-i4). (Scale bar – All CR-stained image scale bars correspond to 50 μm; For SEM of seed sections, scale bar of wheat corresponds to 20μm, while for mungbean, scale bar corresponds to 10μm, for isolated SSPB scale bars correspond to 1μm)

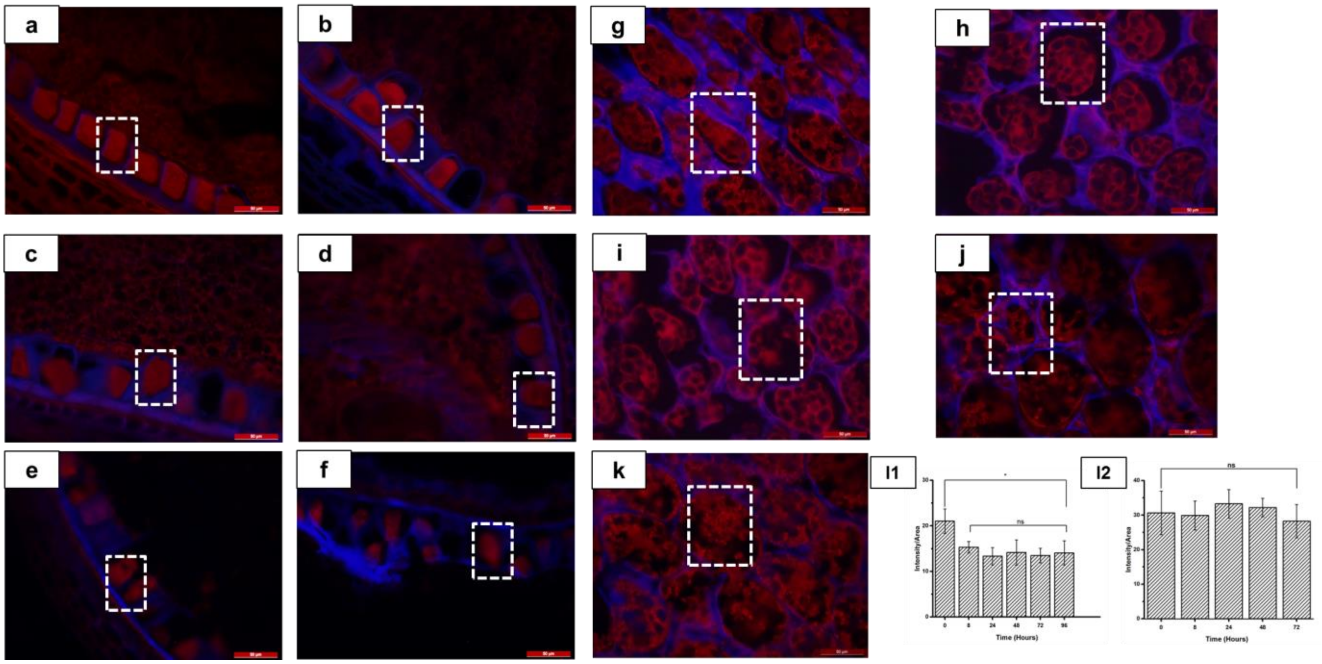


**Fig. S6 Quantitation of the proteinaceous signal in the germinated seed sections.**

Acid fuchsin and calcofluor white staining of wheat seeds are represented at 0 (a), 8 (b), 24 (c), 48 (d), 72 (e) and 96 (f) hours, whereas mungbean seeds are represented at 0 (g), 8 (h), 24 (i), 48 (j) and 72 (k) hours. Quantitative comparison for wheat (l1) and for mungbean (l2), represent the quantitation of the protein signals at each time point. In wheat, the protein signal decreases in the first 8 hours but then becomes constant, whereas in mungbean, the overall protein content remains same till 72 hours of germination. (Gamma value for each image is 1.3-1.4) (t-test, *p=0.02-0.03) (Scale bar – All scale bars correspond to 50 μm)

**a**

| Time (Hours) | Monocots | | | Dicots | | |
|---|---|---|---|---|---|---|
| | Amyloid-like | Amyloid | Protein | Amyloid-like | Amyloid | Protein |
| 0 | +++ | +++ | +++ | +++ | +++ | +++ |
| 8 | + | +++ | ++ | +++ | +++ | +++ |
| 24 | + | +++ | ++ | ++ | +++ | +++ |
| 48 | - | +++ | ++ | + | +++ | +++ |
| 72 | - | - | ++ | + | +++ | +++ |
| 96 | - | - | ++ | Data could not be obtained | Data could not be obtained | Data could not be obtained |

**b**

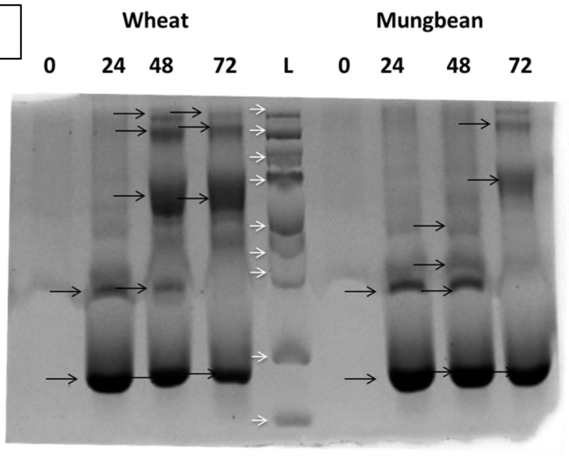


**c**

| Protein body (0 HOUR) | Protein body (48 HOUR) | Protein body (0 HOUR) | Protein body (48 HOUR) |
|---|---|---|---|
| Globulin 3A | Globulin 3A | 8S Globulin beta isoform | 8S Globulin beta isoform |
| HMW glutenin | Globulin 1 | 8S globulin alpha | 8S globulin alpha |
| Globulin 1 | Em protein | Basic 7S globulin-2-like | Oleosin 1 |
| Beta amylase | Triticin | Mungbean seed albumin | Aminopeptidase |

**Fig. S7 The presence of amyloid-like, amyloids and protein content at different time-points of germination.** The time-dependent presence of amyloid and amyloid-like content during germination (a), SDS-PAGE pattern of the disrupted SSPB fibrils at each time-point of germination [Ladder from top– 124, 91,71,54,43,33,29,16 and 10 kDa] (b) and MS-MS analysis of these protein bodies.

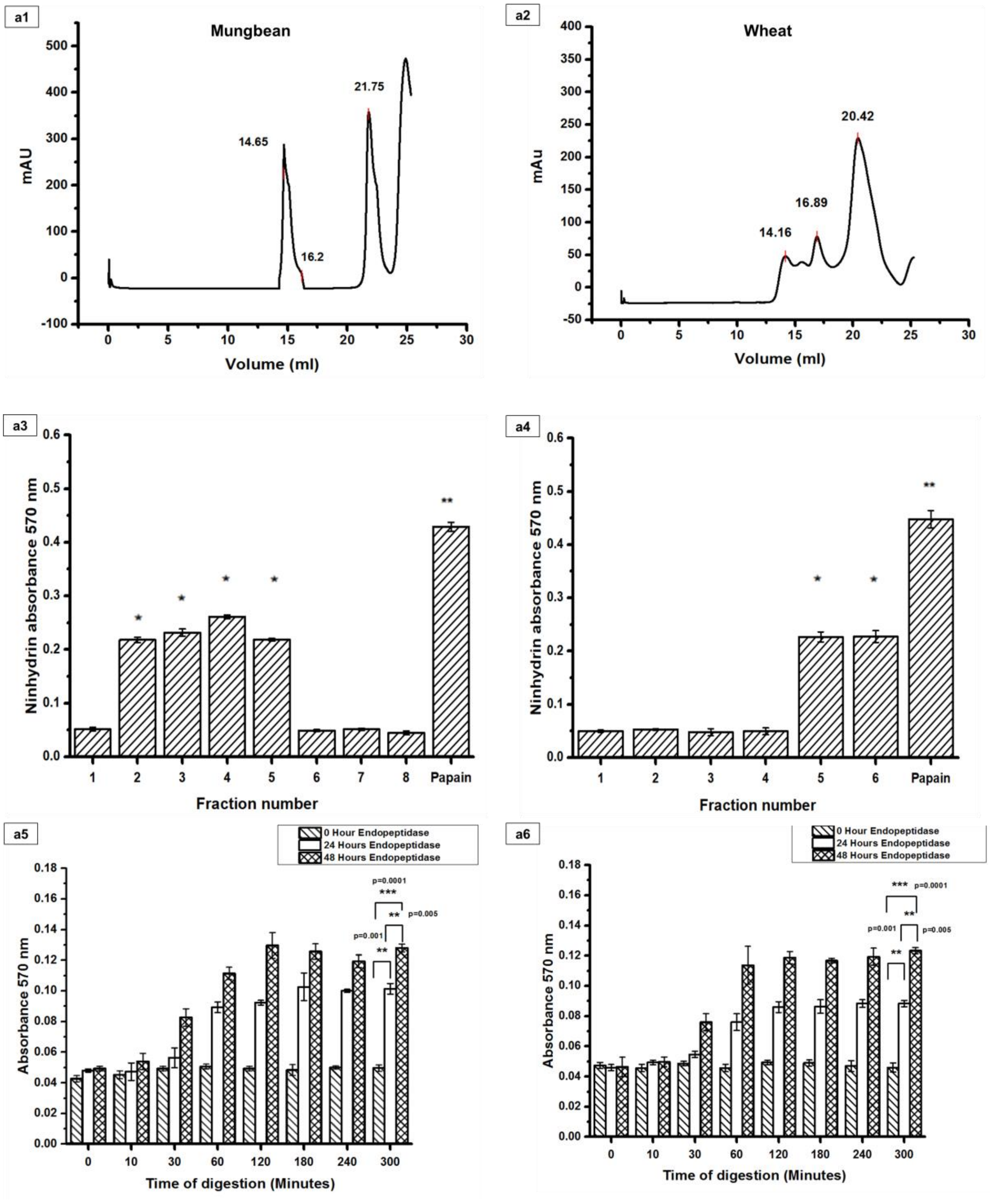


**Fig. S8. Seed endopeptidase characterization and *in-vitro* biological activity**

Representative SEC spectra of the water-soluble fraction of mungbean and wheat (a1-a2) Enzymatic activity of the protein-containing fractions collected from mungbean (a3) and wheat (a4) (*p=0.02-0.04; **p=0.006-0.008) and the time-dependent protease activity of mungbean (a5) and wheat (a6). The fractions with enzymatic activity were incubated with casein as a substrate and the amino acids released are quantified by ninhydrin assay.

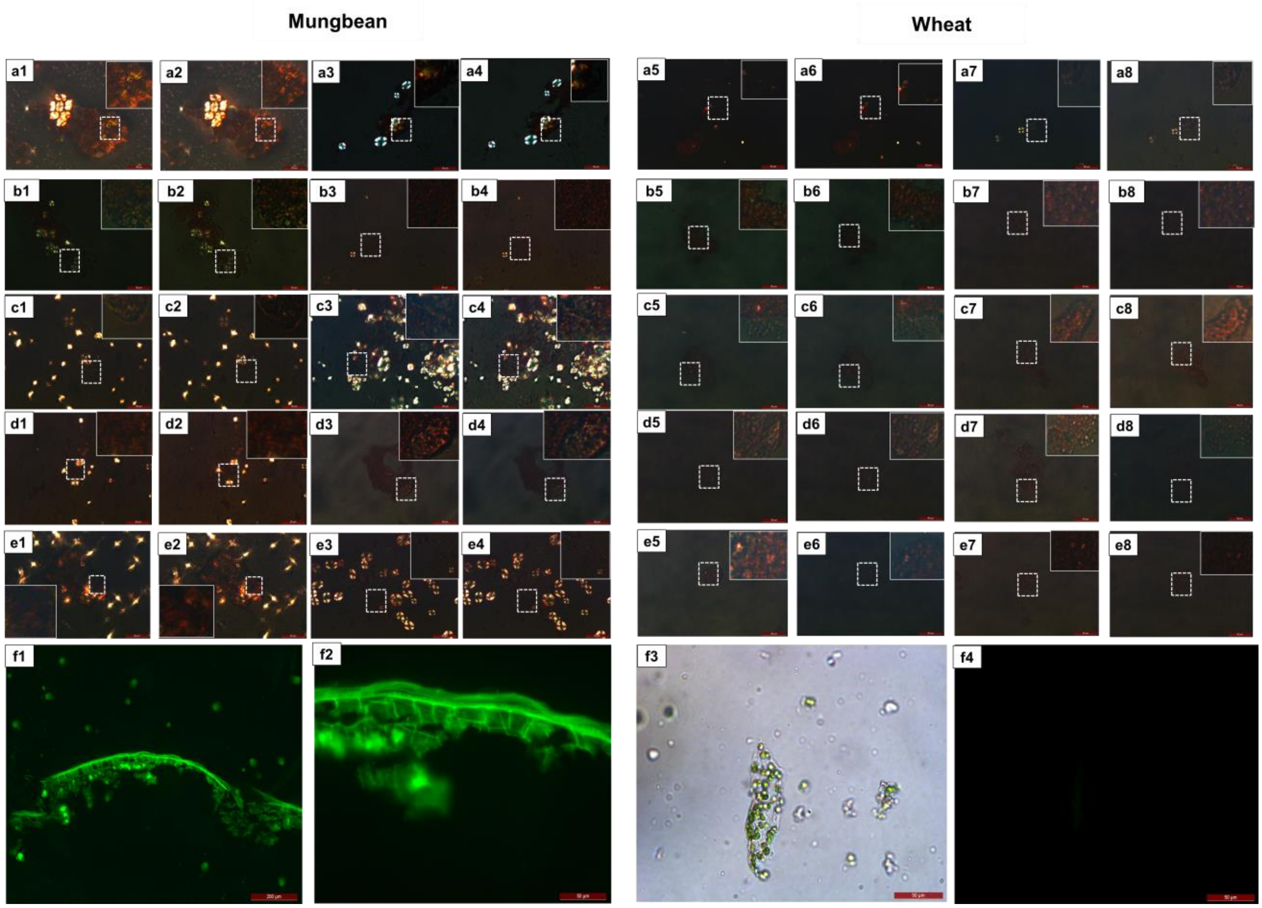


**Fig. S9. Congo red staining of protoplasts treated with exogenous molecules and protoplast control for isolation and staining**

Mungbean (a1-a2) and wheat (a5-a6) protoplasts treated with buffer at 0 hour and after a treatment of 2 hours (a3-a4, a7-a8), shows amyloid signatures are still present. Mungbean and wheat protoplasts treated with seed endopeptidase at 0 hour (b1-b2, b5-b6) and 2 hours (b3-b4, b7-b8), exhibits that birefringence of amyloid structures is not visible. Protoplasts (c1-c2, c5-c6) treated with endopeptidase and PMSF at 0 hours and 2 hours (c3-c4, c7-c8), show that amyloid signatures are retained. Similar retention of amyloids is observed in case of protoplasts treated with GA and PMSF (d1-d2, d5-d6) at 0 hours and 2 hours (d3-d4, d7-d8). However, treatment with ABA and endopeptidase (e1-e2, e5-e6), lose their amyloid signatures after 2 hours (e3-e4, e7-e8). The dashed white boxes represent areas showing birefringence and solid lined boxes represent magnified portions of the insets. (Scale bars correspond to 50 μm) To ensure that the protoplasts are isolated from the wheat aleurone, the dissected aleurone layer was sectioned and visualized by ThT-staining. (f1-f2) As tissue control of the protoplasts, leaf protoplasts were isolated and stained with ThT and showed non-significant ThT signal. (f3-f4)

**Table S2: List of the number of peptides for each protein identified for mungbean and wheat:**

| Sl No. | Protein | No. of peptides | Protein | No. of peptides |
|---|---|---|---|---|
| 1 | 8S Globulin beta isoform | 41 | Globulin 3A | 119 |
| 2 | Beta conglycinin beta chain like | 16 | Globulin 3B | 46 |
| 3 | 8S globulin alpha isoform | 41 | Globulin 1 | 20 |
| 4 | Beta conglycinin alpha chain | 50 | Gamma Gliadin | 13 |
| 5 | Histone | 3 | Ubiquitin | 18 |
| 6 | Glyceraldehyde-3-phosphate dehydrogenase | 9 | Genome assembly protein | 4 |
| 7 | Mungbean seed albumin | 6 | Actin | 6 |
| 8 | Late embryogenesis abundant protein | 24 | Glyceraldyde-3-phosphate dehydrogenase | 5 |
| 9 | Vicilin like seed storage protein | 31 | Em protein | 11 |
| 10 | Beta conglycinin beta chain like precursor | 33 | Histone H4 | 5 |
| 11 | Glycinin G4 | 58 | Actin | 7 |
| 12 | Actin | 16 | Histone H2b | 12 |
| 13 | Elongation factor 1 | 9 | Reversed protein kinase domain containing protein | 2 |
| 14 | Formate dehydrogenase | 5 | HMW glutenin | 29 |
| 15 | Basic 7S globulin-2 like | 6 | Beta amylase | 35 |
| 16 | Class I heat-shock protein | 17 | Aldo_ket_red domain containing protein | 9 |
| 17 | Alcohol dehydrogenase | 4 | 0.19 dimeric alpha amylase inhibitor protein fragment | 11 |
| 18 | | | AAI domain containing protein | 4 |
